# Use of viral motif mimicry improves the proteome-wide discovery of human linear motifs

**DOI:** 10.1101/2021.06.25.449930

**Authors:** Bishoy Wadie, Vitalii Kleshchevnikov, Elissavet Sandaltzopoulou, Caroline Benz, Evangelia Petsalaki

## Abstract

Linear motifs have an integral role in dynamic cell functions including cell signalling, the cell cycle and others. However, due to their small size, low complexity, degenerate nature, and frequent mutations, identifying novel functional motifs is a challenging task. Viral proteins rely extensively on the molecular mimicry of cellular linear motifs for modifying cell signalling and other processes in ways that favour viral infection. This study aims to discover human linear motifs convergently evolved also in disordered regions of viral proteins, under the hypothesis that these will result in enrichment in functional motif instances. We systematically apply computational motif prediction, combined with implementation of several functional and structural filters to the most recent publicly available human-viral and human-human protein interaction network. By limiting the search space to the sequences of viral proteins, we observed an increase in the sensitivity of motif prediction, as well as improved enrichment in known instances compared to the same analysis using only human protein interactions. We identified > 8,400 motif instances at various confidence levels, 105 of which were supported by all functional and structural filters applied. Overall, we provide a pipeline to improve the identification of functional linear motifs from interactomics datasets and a comprehensive catalogue of putative human motifs that can contribute to our understanding of the human domain-linear motif code and the mechanisms of viral interference with this.

## Introduction

Short Linear Motifs (SLiMs; also, often referred to as ELMs - Eukaryotic Linear Motifs) are short linear peptides approximately 3-10 amino acids long, that have a specific sequence pattern that is recognized by interacting domains (Van Roey et al., 2014). They typically lie in disordered regions of proteins, and they are mediators of critical, usually transient interactions involved in dynamic cell processes such as cell signaling, protein degradation, the cell cycle and others (Davey et al., 2011). Therefore, mutations in such motifs often lead to disease, and indeed they have been found to be enriched in cancer (Mészáros et al., 2017; Reimand et al., 2015; Uyar et al., 2014), and several other diseases (Ahmed et al., 2003; Furuhashi et al., 2005; Meyer et al., 2018; Pandit et al., 2007). Gene fusions in cancer often remove SLiM-containing protein regions and the capacity for gene regulation (Latysheva et al., 2016). Due to their short length, small number of specificity-determining residues and disordered nature, they are characterized by high evolutionary plasticity (Davey et al., 2015). They have therefore been used throughout evolution to rewire biological networks, increase their complexity and can often arise through convergent evolution (E. Davey et al., 2012).

A prime example of convergent evolution for the formation of linear motifs are those present in viruses and other pathogens (Davey et al., 2011; Via et al., 2015). Viruses often use molecular mimicry to target hubs or other strategic nodes in host networks and hijack cellular functions. For example the HIV-1 Nef protein interacts with the SH3 domain of the Src family kinases Hck, Lyn and c-Src via a PxxP motif, which leads to their strong activation (Trible et al., 2006). It has been shown that viruses often act in similar ways as oncogenic variations to cause cancer (Rozenblatt-Rosen et al., 2012). A well-studied example of this is the LxCxE motif in the E7 protein of HPV and other viral proteins (Felsani et al., 2006) which acts to inactivate the retinoblastoma protein leading to deregulation of cell cycle control; this inactivation is also common in non-viral cancers (Horowitz et al., 1989; Weinberg, 1995). Viral motifs are generally similar to the host ones, however they tend to be of higher affinity so as to outcompete them (Sheng et al., 2006).

Due to their small size and often low complexity, linear motifs tend to be difficult to discover both by experimental and computational methodologies. In the past decade the availability of high-throughput protein interaction and genomic datasets (Huttlin et al., 2017; Luck et al., 2017; Orchard et al., 2014) have allowed significant advances to be made in developing computational tools for motif discovery. DiliMot (Neduva et al., 2005) and SliMFinder (Edwards et al., 2007) are the first methods able to predict the presence of human motifs at scale, by evaluating over-representation of specific motifs in the interactors of a specific protein and combining it with conservation measures. QSLiMFinder (Palopoli et al., 2015) provides additional functionality by requiring the hits to be present in additional query proteins. FIRE-Pro has also been used to discover yeast motifs in proteome scale datasets based on an information theoretic framework (Lieber et al., 2010). The ELM database and server (Kumar et al., 2020), and SLiMSearch (Krystkowiak and Davey, 2017) can search individual protein sequences or entire proteomes respectively for a given motif pattern. Edwards and colleagues (J. Edwards et al., 2012) demonstrated that using these tools to scan the entire human interactome allows the identification of not only new instances of known motifs, but also entirely novel ones. There are multiple other methods available for discovery of *de-novo*, known and user-defined motifs which can be used for simple protein search of motif patterns or at a much larger scale (Edwards and Palopoli, 2015; Hraber et al., 2020).

By scanning the disordered proteome of a large number of viruses for instances of ELMs, several instances of known SLiMs have been identified (Becerra et al., 2017; Hagai et al., 2014; Liu-Wei et al., 2021). And during the current pandemic, multiple resources have emerged to delineate SLiM based viral-host interactions and discover SLiM instances in SARS-CoV-2 (Li et al., 2021; Mészáros et al., 2021; Yang and Shi, 2021). However, this kind of approach doesn’t consider the wealth of information included in the interacting partners of viral proteins. It also doesn’t allow the discovery of motifs that are not already annotated in public databases. Zheng et al. (Zheng et al., 2014) used the available host-viral interactomes to identify domain-domain interactions between them but did not search for enrichment of linear motif-mediated interactions.

Despite advances in computational discovery of linear motifs the problem remains that only a small fraction of identified motifs is likely to be functional. Thus, additional layers of evidence for each hit can be important to help prioritize them. Here, we hypothesize that requiring human motifs to be present also in viral proteins with common interactors will improve their identification, while also pointing to nodes of viral interference with human host networks. We have thus combined proteome-wide motif search methods with domain enrichment, structural bioinformatics and integration with pathogenic variant information, to perform the first systematic search for human linear motifs also present in common viral interactors, based on the latest available human-viral interactome.

## Results

### A workflow for the discovery of functional linear motifs

Since viruses commonly interfere with human protein interaction networks through viral motif mimicry, we hypothesized that requiring putative new linear motifs to be present both in human and viral protein interactors of specific protein domains would result in enrichment of functional motif instances. Our workflow is based on this principle and in addition filters motifs according to other relevant parameters, such as domain enrichment, predicted binding on domain structure and others (**Figure 1**; Methods).

**Figure 1:**
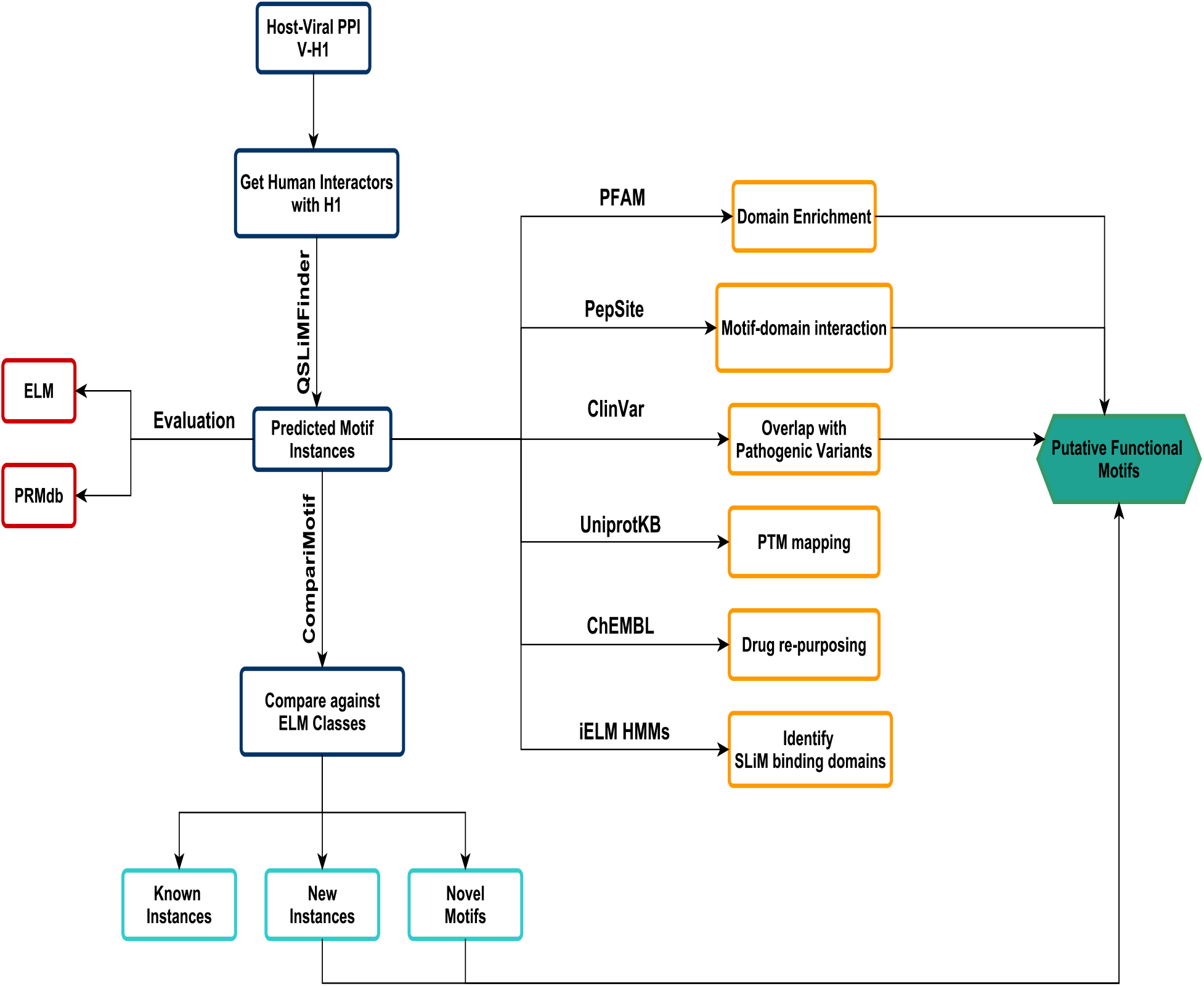
Workflow for discovery of short linear motifs. A schematic describing the overall pipeline for this study. For each from viral-human (V-H1) protein interaction downloaded from IntAct, the respective set of the human interacting partners of H1 were submitted to QSLiMFinder for motif prediction restricting hits to those present also in the respective viral interactor. Then, the predicted hits were evaluated against both ELM and PRMdb to measure how well they capture known hits. In addition, they were also compared against known ELM classes to further differentiate between known, new and novel motif instances. Finally, they were subjected to multiple downstream filters to reduce false positives and further enrich functional instances. Arrow labels show the method/ resource used for each step. V: Viral Protein; H: Human protein which directly interacts with viral protein.

In brief, for each known human-viral protein interaction pair (H1-V), we first used QSLiMFinder (Palopoli et al., 2015) to identify sequence patterns that were both present in the respective viral protein and enriched in the protein interacting partners of the human protein H1 of the query pair (H1-H2, H1-H3 etc). For brevity, throughout the text, V denotes a viral protein that potentially carries a motif, H2, H3 etc human putative motif-carrying proteins and H1 the human domain-carrying protein that interacts with both V and the H2, H3 etc proteins. We thus identified 8,403 putative linear motif instances on 2,757 unique proteins (2457 human and 300 viral) **(Supplementary Table 1)**. An interaction-mediating motif is typically recognized by specific domains. To reduce the potential spurious motifs from our identified set, we thus imposed as an additional filter a requirement for a domain enrichment in the interacting partners of each motif-bearing protein (adjusted p-value <0.05 & present in at least 5 interacting partners **(Supplementary Table 1**). This reduced our initial list to 1,970 motif instances presenting 1.8-fold higher enrichment in proteins including true SLiMs compared to raw predicted hits (**Figure 2 C,D; Methods**). As part of the workflow, we also use HMM profiles from the iELM web server (Weatheritt et al., 2012) to identify domains that have a match to domain profiles known to bind to known ELM motifs. While this leads to an improvement in the enrichment of known motifs (**Figure S1**), and is provided for information **(Supplementary Table 1**), we don’t use it as a filter, as there are only available domain-specific HMMs for a small fraction of our H1 interacting proteins (125/517).

**Figure 2:**
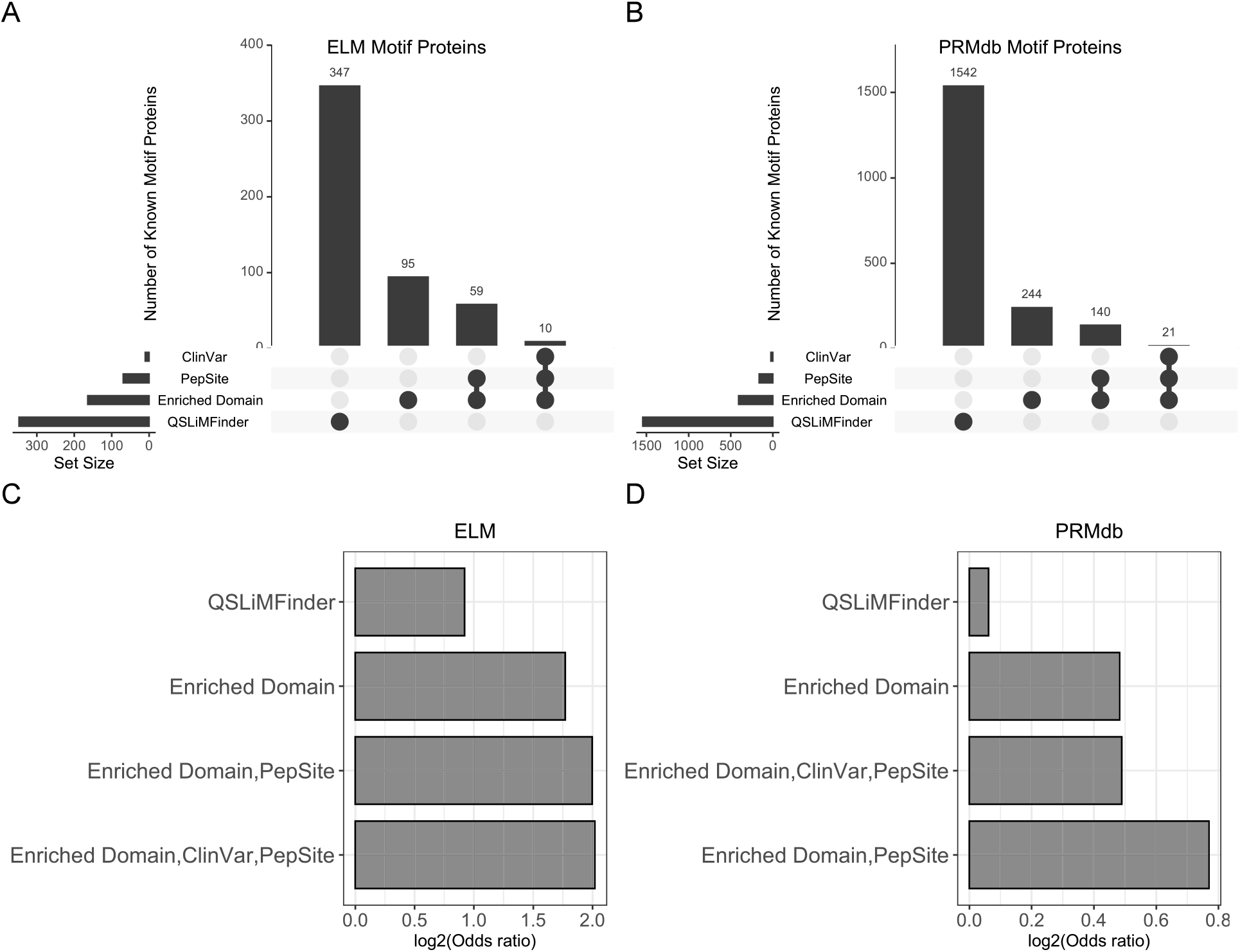
Evaluation of known motif-carrying proteins. (A & B) UpSet plots showing the overlapping number of known motif-carrying proteins that are found in either ELM (A) or PRMdb (B) and predicted datasets defined by a single or multiple filters. (C & D) Bar graph plotting the enrichment of known ELM (C) or PRMdb (D) motif-carrying proteins represented by log2(odds ratio) for each set of applied filters. Odds ratio was calculated based on a one-tailed fisher-exact test using query input to QSLiMFinder as background. All filters mentioned were applied on the raw output of the QSLiMFinder method (See also **Figure S1**)to mean random DMI of 17.29 and 16.5 with an enrichment score of 5 and 2.7 (FDR 0.2 vs FDR 0.37) for host-viral and human-only hits, respectively (**Figure S2 C,D**). Therefore, the observed DMIs are significantly higher than expected for both datasets (P < 0.001) with the host-viral screen having ∼2X fold higher enrichment than the human-only one.

Using a structural bioinformatics strategy, we then sought to further reduce false positive hits. Specifically, we used PepSite2 (Petsalaki et al., 2009; Trabuco et al., 2012), which identifies binding pockets for peptides on protein structures, and tested whether the domains we identified above indeed carry a pocket that can accommodate the peptides carrying our putative new motifs. This step was dependent on the availability of the relevant protein domain structure, and thus was applicable to 7,941/8,403 of our identified motif instances. The result was 1,033 high quality putative linear motif instances for which we found evidence at the sequence, network and structural level and showed a further enrichment in known motif-carrying proteins **(Supplementary Table 1; Figure 2 C,D)**.

Finally, to provide functional support for our predicted motifs and further study the crosstalk between viral SLiMs and disease pathways, we mapped the predicted motifs to pathogenic variants from ClinVar (Landrum et al., 2018). We found that ∼13% of our predicted motif instances were associated with a ClinVar variant, with 27% of these mapping to a pathogenic variant and ∼85% having a significant fit on the structure, as predicted by PepSite2, to a pocket of a relevant domain (OR = 3.47, p-value = 5.3×10^−18^). We thus identified 105 ‘gold’ functional linear motif instances that present a 2.15-fold enrichment in proteins including true SLiMs compared to the initial QSLiMFinder scan **(Supplementary Table 1; Figure 2 C,D).**

To evaluate the benefit of requiring motifs to be present in both human and viral interaction partners of a protein, this workflow was repeated as described also on the human interactome without this requirement (**Supplementary Table 2).** Starting from 3,994 initial predicted motifs from SLiMFinder (Edwards et al., 2007), our filtering process resulted in 29 ‘gold’ human motifs. We found that the inclusion of the viral motif requirement as a filter resulted in a 2.1 fold higher number of motifs and in improvement in almost all metrics of performance (**Figure S2 A,B**; 8,403 vs 3,994). We additionally used SLiMEnrich (Idrees et al., 2018), to evaluate the level of enrichment of known Domain-motif interactions (DMIs) compared to random predictions for both analyses. Using ELM as background, 86 and 45 non-redundant DMIs were predicted compared

### Similarity between discovered motifs and known motif classes

To further classify the predicted motifs, we compared their regular expression patterns to those of known ELM identifiers using CompariMotif (Edwards et al., 2008), which scores each comparison based on the shared information content between two motifs. We found that the ELM classes were equally represented, with a slightly higher representation of Ligand binding sites (**Figure S3A**). When we only consider high-similarity comparisons (i.e having CompariMotif scores >= 3) and excluding CLV (cleavage) and TRG (targeting) classes (**Figure 3; Methods**), we found that 18 out of 77 motif patterns were part of previously known ELM instances, whereas the rest were novel ELM instances matching to 30 ELM identifiers. The top matched ELM identifiers are LIG_PDZ_Class_1, LIG_ULM_U2AF65_1, and DOC_MAPK_gen_1 showing high similarity with 30 out of 77 predicted motif patterns (**Figures 3, S3B**). The aforementioned ELM classes have been previously reported to be involved in multiple host-viral interactions throughout the viral life cycle (James and Roberts, 2016; Pabis et al., 2019; Evans et al., 2010). If we only consider the ‘gold’ instances, approximately 50% of the predicted motif patterns are associated with CFTR protein which bears multiple disease mutations, mapped through various host-viral interactions (**Figure 4A & B**).

**Figure 3:**
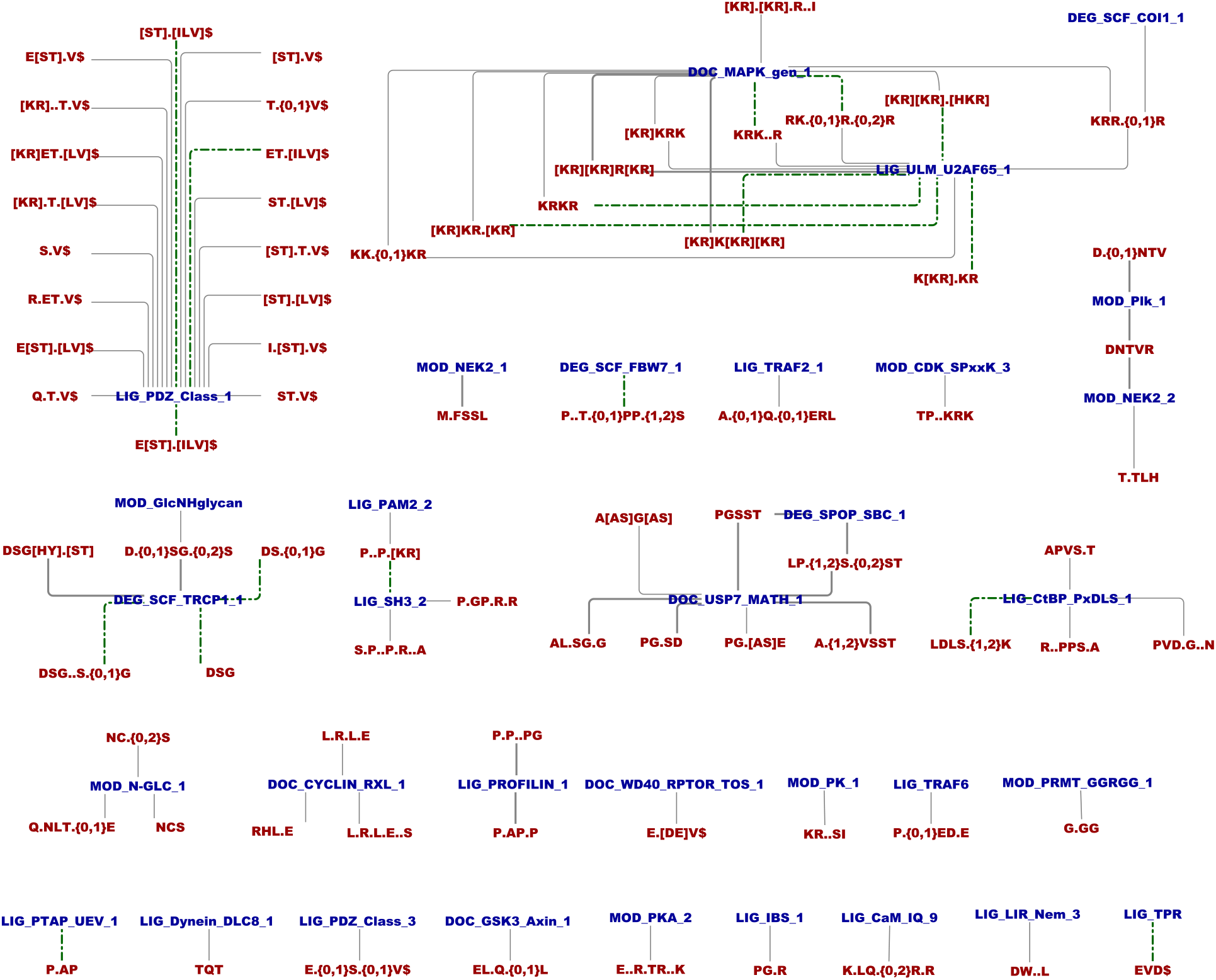
Comparison of predicted motif patterns to known ELM classes. Network showing the connection between predicted motif patterns with > 60% similarity to known ELM classes according to CompariMotif scores. For the sake of visualization, CLV and TRG classes were excluded as they tend to be easily discovered because of their shared location, and thus multiple patterns can be promiscuously mapped to few ELM classes. Predicted motif patterns are shown in red, while ELM classes are shown in blue. Green dash-dotted edges represent motif patterns that match to a known ELM instance, while solid edges represent new instances.

**Figure 4:**
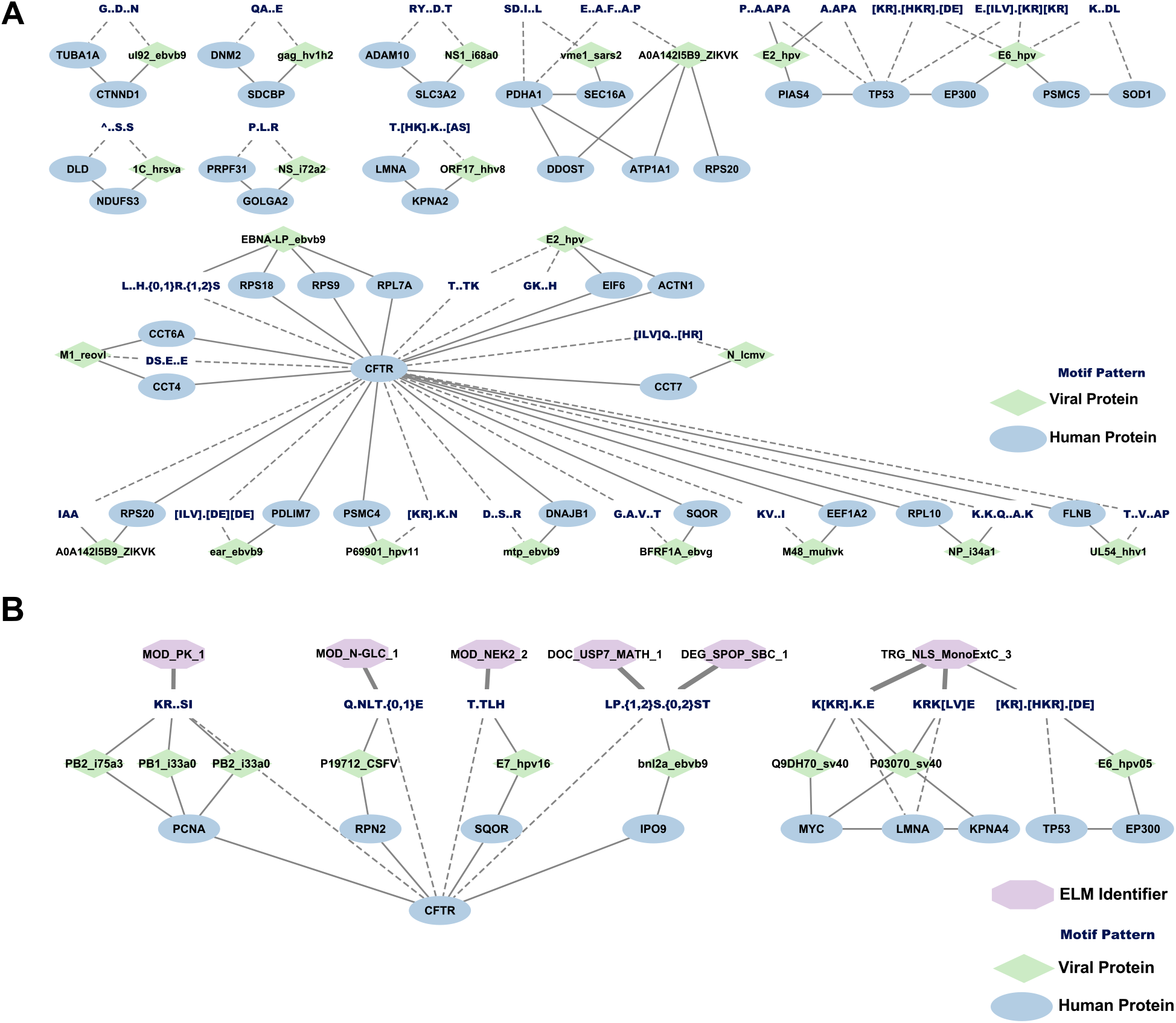
Putative functional motif instances. Network showing novel motif instances (A) and new ELM instances (B) associated with a ClinVar pathogenic variant in addition to confirmed structural binding with enriched domains according to PepSite. Stricter significance cutoffs were used for this figure compared to the ‘gold set’ (QSLiMFinder p-value < 0.05; PepSite p-value < 0.05) for simplicity of visualization. Nodes colored in green, cyan and dark blue represent viral, human and predicted motif patterns, respectively. Viral protein labels include protein name and viral strain separated by “_”. Dashed edges are only drawn between motif patterns and human proteins, where a ClinVar variant is associated with that motif pattern that is predicted on the respective human protein. Solid edges between proteins represent PPI as retrieved from the IntAct database.

### Functional enrichment, essentiality and pathway-specificity of linear motifs

Next, we sought to identify the functional pathways in which our predicted DMIs are involved. First, we performed Gene Ontology (biological process - GOBP) enrichment (Ashburner et al., 2000; Gene Ontology Consortium, 2021) for H1 proteins to find overrepresented pathways in each filter combination using all 517 H1 proteins as background (Methods). The top enriched GO terms can be summarized in 4 major categories: regulation of gene expression and nucleosome assembly, metabolic processes and protein-modification, interaction with host and symbiotic processes, and nuclear import/export and protein localization (**Figure 5**). Some filter combinations (e.g. Enriched Domains, ClinVar and PepSite) are almost exclusively enriched in protein transport and localization (**Figure 5**). Additionally, we performed GOBP enrichment analysis of either H1 domains or domains that interact with predicted human motifs, using the pfam2go (biological process - PFAM2GOBP) associations (see Methods) as background (Mitchell et al., 2015). Only 180 out of possible 679 H1 domains and 1,204 out of possible 4,674 enriched domains are annotated in PFAM2GOBP, thus this analysis was performed only on the subset for which annotations were available. As in H1 GOBP enrichment, we identified overrepresented pathways in each filter combination (Methods). Only 3 filter combinations had significantly enriched terms (adjusted p-value < 0.05): iELM, Enriched Domain & Pepsite, and ClinVar and PepSite. Domains in H1 proteins were mostly enriched in metabolic processes, specifically nitrogen-compound metabolism and RNA synthesis processes (**Figure 6 A,B**). Domains in interacting partners of predicted host motifs were mostly enriched in protein phosphorylation, RNA biosynthetic processes, and protein transport and localization (**Figure S4 A,B**). Moreover, we also measured the semantic similarity between the enriched GO terms for H1 domains and domains that pass either the PepSite filter, the Enriched Domain filter or both. In concordance with the aforementioned enriched terms, the most prominent PFAM domain-associated biological processes represented are transcription-related (Myc_N, Creb_binding, KIX), translation related (Ribosomal_S17, Ribosomal_S13), and phosphorylation-related (Pkinase, PK_Tyr_Ser-Thr, Pkinase_C) (**Figure 6C, S4C**).

**Figure 5:**
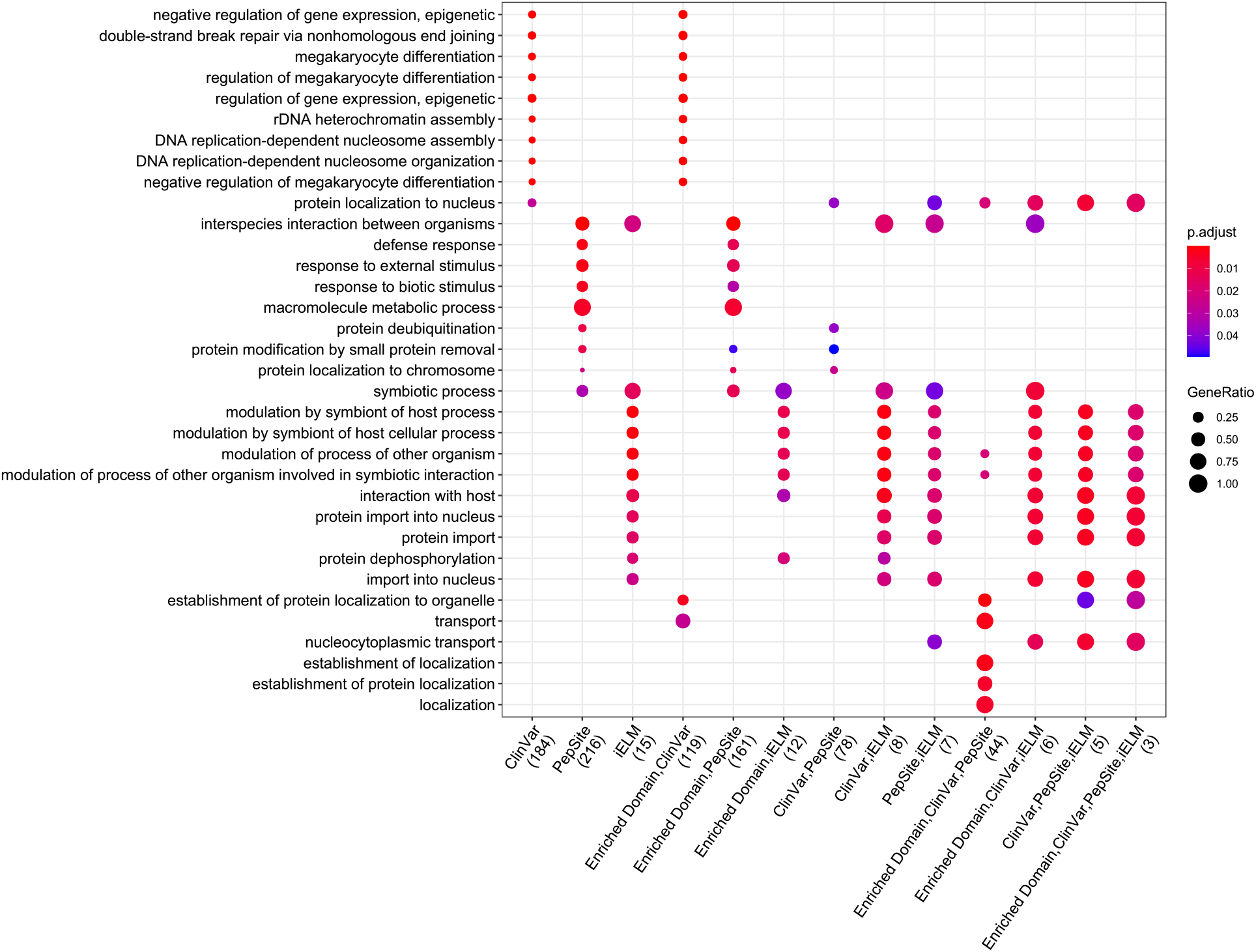
H1 protein GO enriched pathways. Dot plot showing the enriched GO terms (biological process) for each filter combination. This is based on an overrepresentation test using the 517 H1 proteins as background. The color scale corresponds to adjusted p-value and the size of dots corresponds to the proportion of overlapping proteins between input query and annotated proteins for each GO term. The number of enriched H1 proteins in each dataset is enclosed in parentheses for each filter combination annotated under x-axis labels.

**Figure 6:**
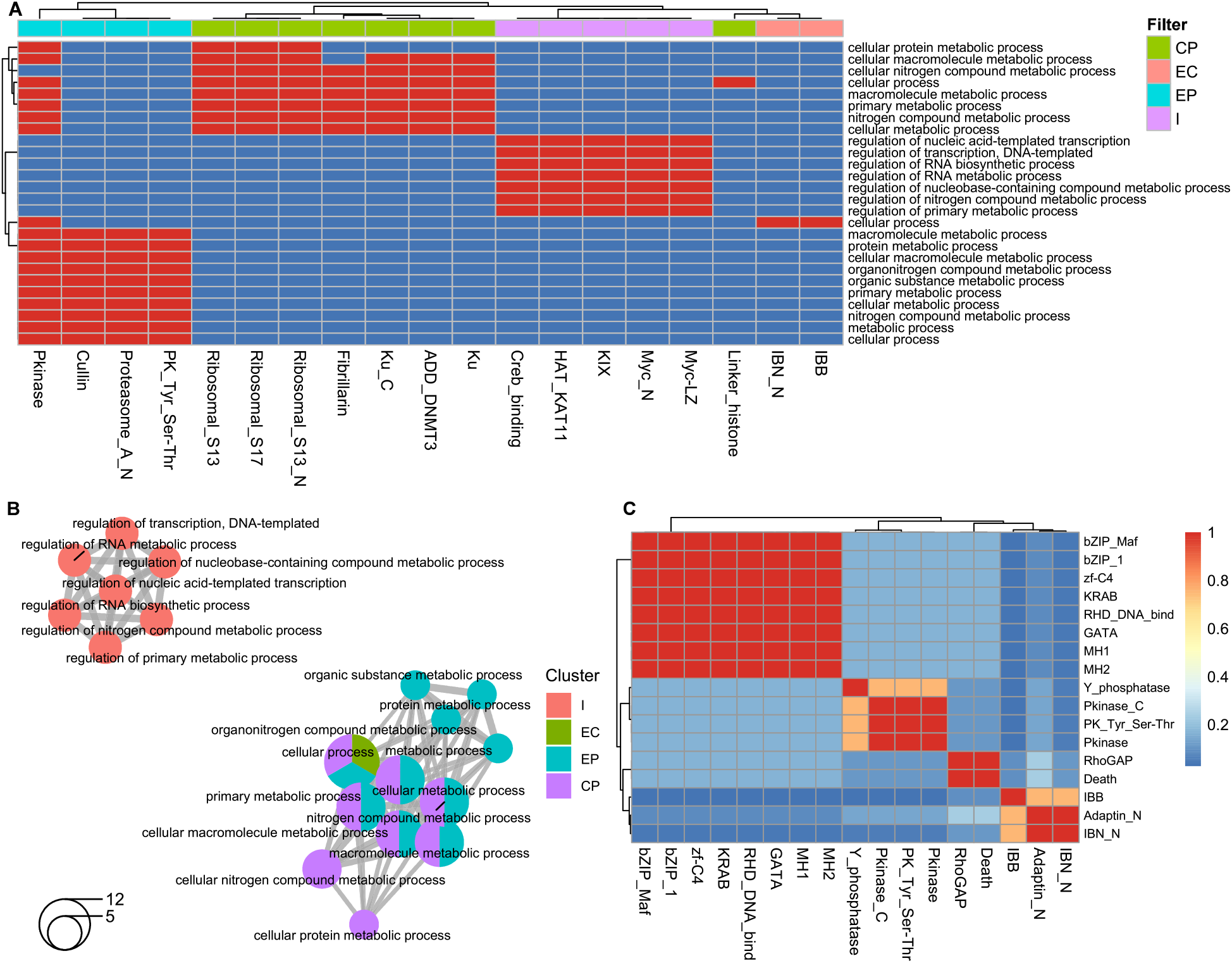
GO pathway enrichment of PFAM domains in H1 proteins. (A) Binary heatmap describing association between enriched GO terms (biological process, y-axis) and the corresponding overlapping domains between input dataset and PFAM2GOBP annotations (x-axis) for a given term, where red and blue boxes represent presence or absence of enriched terms for a given domain, respectively. Only Domain-pathway pairs with adjusted p-value < 0.05 are displayed. Domains are clustered by filter combinations shown as colored annotation bars above columns. E, Enriched Domains; P, PepSite; C,ClinVar; I, iELM. (B) Network showing the Jaccard similarity coefficient between enriched GO terms. Each node is a pie chart colored by filter combinations and the size of the nodes correspond to the number of domains. E, Enriched Domains; P, PepSite; C,ClinVar; I, iELM. (C) Heatmap showing GO semantic similarity between enriched domains calculated using the “Wang’’ method (Wang et al., 2007) between GO terms for each domain-pair. The color scale represents the similarity score where 1 denotes complete overlap, and 0 denotes no similarity..

We then explored whether the domains involved in our predicted domain-motif interactions tend to be specific to certain GOBP or KEGG (Kanehisa and Goto, 2000; Kanehisa et al., 2021) pathways using a published network-based scoring approach (Shim et al., 2019). The top-scoring (z-score > 1) processes and pathways associated with at least 3 filter-combinations are: cell motility, translation, and regulation of transcription (**Figure S5A**), and DNA repair and transcriptional misregulation in cancer (**Figure S5B**), respectively.

To gain more insight into the types of proteins that are involved in motif-domain interactions we also assessed the essentiality of both H1 proteins and predicted motif-carrying proteins. For our H1 proteins, all filter combinations including Enriched Domains, PepSite and ClinVar were significantly enriched in essential genes. Combinations including iELM datasets were mainly enriched in context-essential genes (**Figure S6**), although not significant due to the very low numbers of H1 and predicted motif-proteins covered.

### Identification of candidates for Drug repurposing

Given that for all our human predicted motifs there was a corresponding viral motif targeting the same domain-carrying protein (H1), we sought therapeutic drugs that target the latter. Our hypothesis was that, if those drugs are indicated for the ClinVar diseases resulting from a pathogenic mutation in the relevant predicted motifs, they may be suitable for repurposing for the corresponding viral infection.

By mining the ChEMBL database (Gaulton et al., 2017; Mendez et al., 2019) we retrieved 13 molecules targeting 5 H1 proteins that also had drug indications matching the relevant ClinVar disease (**Figure 7; Supplementary Table 3**). Roscovitine (Seliciclib), which targets PSMC4 (26S proteasome regulatory subunit 6B), was one of the identified compounds. PSMC4 interacts with CFTR, where the pathogenic mutation (AlA412del) was found in the motif pattern ([KR].K.N). The same motif pattern was also observed in HPV11-E5B (Human Papillomavirus 11 – E5B) which is associated with PSMC4 through Tandem Affinity Purification (TAP) / MS (Rozenblatt-Rosen et al., 2012). Roscovitine has been previously reported to prevent the proteasomal degradation of F508del-CFTR through a CDK-independent mechanism, thus restoring the cell surface expression of CFTR in human CF (cystic fibrosis) airway epithelial cells (Norez et al., 2014). On the other hand, there have been several reports linking HPV E5 proteins with the host ubiquitin-proteasome system to modify and regulate host proteins (Wilson, 2014). Accordingly, Roscovitine can be potentially repurposed as pharmacological therapy for HPV 11 infections. Moreover, several other drugs that we identified (**Figure 7**) already have indications in ChEMBL or evidence for reducing the relevant viral infection, such as Sirolimus and Everolimus for Influenza A infection (Alsuwaidi et al., 2017; Murray et al., 2012), Vorinostat and Cytarabine for human herpesvirus type-8 (hhv8) (Hogan et al., 2018; Jc et al., 2020), and Erlotinib, Vatalanib, and Dasatinib for Hepatitis C (HCV) (Lupberger et al., 2011; Elsebai et al., 2016).

**Figure 7:**
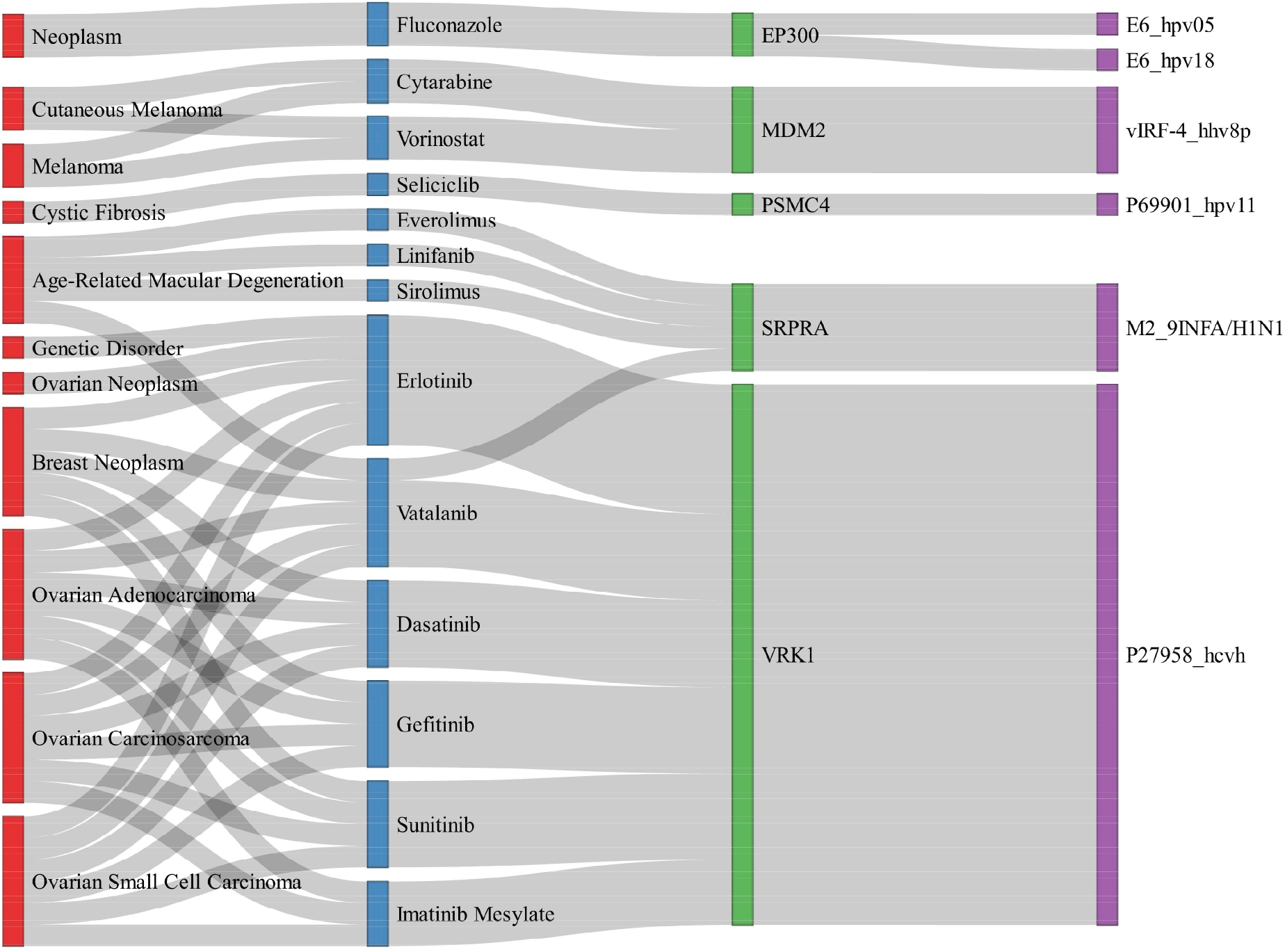
Potential Drug candidates for repurposing. Sankey network displaying the relationship between ClinVar diseases and their indicated ChEMBL drugs that can be potentially repurposed for the respective viral infection. Drugs target the host domain protein (H1) which directly interacts with the relevant viral protein (V1). Nodes colored in red, blue, green and violet represent ClinVar disease, ChEMBL drugs, host protein, and viral protein respectively.

## Discussion

Despite their genome size constraints, viruses manage to hijack a myriad of cellular processes and signalling pathways, often by means of SLiM-mediated interactions with the host proteome, thereby taking control of the host cellular machinery to their reproductive advantage (Davey et al., 2011). Moreover, because of their small size, degenerate nature, low complexity, and presence in intrinsically disordered regions, SLiMs are rapidly evolving *ex nihilo* (evolution from “nothing”) due to frequent mutations in these disordered regions, thus allowing a diverse repertoire of motifs to emerge in unrelated proteins through convergent evolution (Davey et al., 2015). Altogether, the nature of these short motifs explains the highly inflated false positive rates that can be computationally discovered throughout the proteome, due to their relatively poor signal to noise ratio (Gibson et al., 2015). While there has been a considerable effort to elucidate the molecular mechanisms by which viral proteins interact with the host proteome, there are relatively few resources (Hraber et al., 2020) highlighting SLiM-mediated interactions, the best of which is currently the ELM database (Kumar et al., 2020). In addition, resources and approaches to discover new linear motifs in general, rely on the largely manual identification of limited new motif instances or the re-discovery of known ELMs in different protein sequences, and in doing so, they fail to consider the wealth of information included in the interacting partners of viral proteins.

Here we describe a proteome-wide approach to identify novel functional human motifs by taking advantage of the latest available human-viral interactome (Orchard et al., 2014), and including evidence from domain enrichment, structural modelling of SLiM-domain interactions (Petsalaki et al., 2009; Trabuco et al., 2012), and mapping ClinVar (Landrum et al., 2018) pathogenic variants on our predicted motifs, to improve both the sensitivity and specificity of computational motif discovery. Overall, using the principle of viral motif convergent evolution, we improved both the sensitivity and the specificity of the motif identification (**Figure S2**) and discovered 74 and 3,673 known and new motif instances, respectively, in addition to 4,656 novel motifs.

Functional analysis of our putative motif-domain interactions showed enrichment in processes known to be mediated by motifs, and importantly also hijacked by viruses. These include protein translocation (Cook and Cristea, 2019), transcriptional regulation (Kropp et al., 2014; Tarakhovsky and Prinjha, 2018), cell signalling (Alto and Orth, 2012), and cellular metabolism (Mayer et al., 2019; Thaker et al., 2019) among others, highlighting that despite a potential high number of false positive motifs identified, our resource is enriched in functional hits that describe biologically relevant processes. These have also been identified in a similar study that performed a proteome-wide analysis of human motif-domain interactions in influenza A viral proteins (García-Pérez et al., 2018). Interestingly we found a significant enrichment of essential genes to be involved in motif-domain interactions. This highlights the importance of such interactions in cell functions and provides additional support for the functionality of our predicted motif-domain interaction set. Our observation can partially be explained by the fact that motif-binding domains and on occasion also motif-carrying proteins can interact with multiple diverse partners, i.e. be hubs, which are known to be more essential than other nodes in protein interaction networks (Puschnik et al., 2017; García-Pérez et al., 2018).

We also highlight specific diseases associated with a disease-causing variant mapped to the shared motif pattern between host-viral and human PPIs, thus providing potential insights into the crosstalk between viral infection and the underlying disease. Interestingly, among the gold set of our hits about 50% putative motif patterns were associated with a pathogenic variant in *CFTR* gene (**Figure 4**). Viral infections have been heavily implicated in the pathogenesis of cystic fibrosis by increasing the susceptibility to other bacterial infections leading to further complications in the respiratory tract in cystic fibrosis patients (Frickmann et al., 2012). It has also been reported that multiple viral proteins with homology to CFTR interact with the same targets as CFTR, to commandeer the CFTR interactome. Several pathways are common between CFTR interactome and the viral life cycle including: endosomal pathways, lysosomal trafficking and protein processing (Carter, 2010). This could explain why we got multiple hits with CFTR and these putative motif-domain interactions shed light on the potential mechanisms by which viral proteins interfere with the CFTR interactome.

In a search for drug repurposing candidates, we also identified drugs that target domain-carrying properties that we predicted to be relevant both for the viral infection and associated ClinVar disease. While only 13 drugs targeting 5 of our proteins were retrieved, several of them already had indications for the relevant viral infection in addition to the disease used to extract the drugs from the database (Alsuwaidi et al., 2017; Elsebai et al., 2016; Hogan et al., 2018; Jc et al., 2020; Lupberger et al., 2011; Murray et al., 2012). This suggests that our collection of paired viral-human motifs could be a good starting point for identifying key nodes to target in the pathogenesis of both the disease associated with the motif mutations and the viral infections.

Although we observed evidence for the presence of functional novel motifs and the recovery of known instances, computational *de novo* SLiM prediction remains a difficult challenge despite the plethora of available SLiM prediction methods, and thus the number of false positive predictions is expected to be high (Prytuliak et al., 2017). Our approach is vulnerable to additional uncertainties, which can be attributed to multiple confounders throughout our workflow in addition to the inherent false positive rate of the motif prediction method. Starting off with the PPI data from IntAct (Orchard et al., 2014), some of the binary interactions reported in IntAct database have weaker interaction evidence and thus an overall low Molecular Interaction score (MIscore) compared to “true” interactions which has been confirmed by several methods and has more literature evidence (Villaveces et al., 2015). Therefore, some of the input host-viral interactions are merely association events as opposed to having strong structural binding evidence and could be introducing noise to the analysis. Furthermore, as with all proteome-wide motif discovery methods, the precision and recall were relatively low when we evaluated the predicted instances against that of ELM’s. Nevertheless, when we used SLiMFinder (Edwards et al., 2007) to apply the same pipeline on all human proteins without restricting the motif space to viral motifs, we obtained even lower precision and recall in most of the evaluations (**Figure S2)**. Therefore, the predicted hits conditioned on the presence of viral motifs recover more known instances with higher accuracy compared to their absence despite their apparent high false positive rate.

Our study provides a considerable number of potential motif instances with several levels of support for functionality that can be prioritized for further investigation and validation studies (e.g. using ITC or Fluorescent Polarization). We show that taking advantage of the principle of convergent evolution allows us to reduce the false positive rate inherent to computational screens for linear motifs, which we then further reduce using additional orthogonal filters. This not only provides an improved catalogue of putative human motifs but also provides indications for points of motif-mediated viral interference that can be easier to target than interactions mediated by larger interfaces. This resource will be valuable towards improved understanding of the biological functions of motif-mediated interactions in humans and viruses. As more human-viral protein interactions, domain structures and other information become available, our workflow will be able to provide evidence for an increasing number of human motifs.

## Acknowledgments

The authors would like to thank EMBL-EBI for funding this project and Ylva Ivarsson for input on the manuscript and validation.

## Author Contributions

Conceptualization: BW, VK, EP, Methodology: BW, VK, EP, Software: BW, VK with contribution from ES, Validation: BW with contribution from CB, Formal Analysis: BW, VK, Investigation: BW, VK, EP, Resources: EP, Data Curation: BW, VK, Writing - Original Draft: BW, EP, with some contribution from VK, Writing - Review & Editing: BW, EP, Visualization: BW, Supervision: EP, Project Administration: EP, Funding Acquisition: EP

## Declaration of interests

The authors declare no competing interests.

## Data and code availability

To ensure reproducibility, all data, code, intermediate and final analysis files, have been made available as follows:

- All, raw and analyzed data that are not already included in Supplementary Tables in this manuscript are publicly available at the Biostudies database with project ID: S-BSST668 (https://www.ebi.ac.uk/biostudies/studies/S-BSST668/)
- All original code is publicly available at https://gitlab.ebi.ac.uk/petsalakilab/HVSlimPred.

## Methods

### Interactome Datasets

We extracted all human-viral interactions from the IntAct database (Orchard et al., 2014) on 18/05/2020 by using the query:“(taxidA:9606 AND taxidB:10239) OR (taxidA:10239 AND taxidB:9606) AND ptypeA:protein AND ptypeB:protein” on 20/05/2020. This resulted in 22,839 binary interactions. We then removed the duplicate entries resulting in a Human-Viral dataset of 15,559 interactions including unique 5,575 proteins from 325 viruses (types and isolates) and humans (**Supplementary Table 4**). Distributions of interactions per virus type are shown in **Supplementary table 5.**

In addition, we extracted all human interactions that had more than 2 types of evidence and at least one of the two interacting proteins was present in the Human-Viral dataset we had already collected. The resulting dataset included 49,000 interactions amongst 10,707 proteins (**Supplementary Table 6**).

### Validation datasets

Our main benchmarking dataset comprises 1,936 known human SLiMs in 1,153 human proteins, extracted from the Eukaryotic Linear Motif (ELM) database on 24.07.2020 (Kumar et al., 2020) (**Supplementary Table 7**). In addition, we considered PRMdb (Peptide Recognition Modules database) which is based on large-scale peptide phage-display methods (Teyra et al., 2020) and comprises 386 PRM modules covering 50 structural families in 12,771 human proteins spanning 47,657 SLiMs (**Supplementary Table 8**).

### Identification of linear motifs in viral-human protein interaction networks

QSLiMFinder (Palopoli et al., 2015) was used for the identification of motifs in our networks with the default settings in terms of filtering except for setting the maximum number of sequences to evaluate to 800 (dismask=T, consmask=T, cloudfix=T, maxseq=800, gnspacc=F). Specifically, for each viral protein (V1) interacting with a specific human protein (H1) we searched for motifs that are enriched in the known human protein interaction partners of H1 and the viral protein. We also imposed the restriction that the motif must also be present in the viral protein.

### Enrichment of human protein domains as potential motif interactors

For each viral-human V-H1 protein interaction we extracted all human interactors of H1 from the IntAct database (Orchard et al., 2014). To avoid bias towards specific domain types due to sequence similarity of these interactors, we then used cd-hit (Fu et al., 2012; Li and Godzik, 2006) to reduce redundancy at 70% sequence identity level. We then used a two-tailed fisher-test (Fisher, 1934) to calculate the enrichment of observing a specific domain for each interaction set compared to the background. These p-values were FDR corrected using the Benjamini-Hochberg approach (Benjamini and Hochberg, 1995). We only considered as enriched the domains that had a significant adjusted p-value in terms of enrichment compared to the background (adjusted p-value < 0.05). As an additional stringency filter, we required that the identified domain was present in at least 5 interacting partners of the motif carrying protein. This choice was made after evaluating multiple different cutoffs for the adjusted p-value and domain numbers and selecting the best performing combination (**Figure S7 – also see next section**).

### Selection of domain enrichment filter

To select possible filters for functional motif enrichment, we first tested the effect of considering enriched domains on the enrichment of known hits. We tried different combinations of parameters concerning the minimum number of interaction partners for a specific domain (1-10), odds ratio (>1 or >4), and adjusted p-value (0.01, 0.05, 0.1) for a total of 60 datasets. Based on the scores for all 3 levels, a minimum of at least 5 interaction partners returned the best results, where the difference was more pronounced in PRMdb (Teyra et al., 2020) compared to ELM (Kumar et al., 2020) (**Figure S7**). Therefore, the predicted hits were filtered so that the identified domain was present in at least 5 interaction partners of the motif carrying protein with an adjusted p-value of < 0.05. The previous dataset was used in further downstream analyses as another functional filter to reduce false positives and will be referenced as “Enriched Domain”. Additionally, we applied the same evaluation pipeline using combinations of functional filters described below including: ClinVar, PepSite, Enriched Domain, and iELM HMMs to check which functional filters are better at capturing known instances.

### Comparison of discovered motifs with known ELM classes

To further classify the predicted motifs, we compared their regular expression patterns to those of known ELM identifiers using CompariMotif (Edwards et al., 2008), which scores each comparison based on the shared information content between two motifs. We grouped the matched relationships identified by CompariMotif into 5 confidence levels ranked by their proposed heuristic score. This considers the number of matched positions and a normalized score corresponding to the information content between the respective motifs. For each of the six major ELM classes, (ligand (LIG), cleavage (CLV), docking (DOC), degradation (DEG), post-translational modification (MOD), and targeting (TRG)), we performed an all Vs all comparison against the relevant ELM identifiers (**Supplementary Table 1**). The predicted instances were also classified into known and new instances based on whether the instance overlaps with a true positive instance in the ELM database or not, respectively (**Figure S3A**). In addition, we also checked the concordance between CompariMotif similarity score and the known instances that we predicted, confirming that indeed high CompariMotif scores were able to identify true ELM class labels (AUC 0.791-0.829 depending on the set evaluated).

### Evaluation of predicted motifs

Fair evaluation of linear motif identification methods is complicated partly due to: a) the difficulty to define terms like ‘true positive’, because motif instances often overlap and are found in duplicates and b) the very unbalanced nature of the validation dataset with very few known motifs and undefined, but likely large, numbers of true but not yet discovered motif instances. Thus to provide performance metrics from different points of view, we performed the evaluation at 3 levels (motif-carrying protein, motif instances, protein-domain interaction) using both ELM (Kumar et al., 2020) and PRMdb (Teyra et al., 2020) as separate validation datasets.

For the protein-level enrichment, we measured the enrichment of true-positives in our predicted dataset using a one-tailed Fisher’s exact test (Fisher, 1934), where the odds ratio represents the magnitude of the enrichment. True positives are the number of motif-carrying proteins present in both the predicted dataset and the validation dataset regardless of whether the predicted protein has the right motif or is found in the right location. For this level, we used the odds ratio to evaluate enrichment of true positives in each filtered dataset, as a measure of improved quality of the predictions after each filter.

For the motif and protein-domain levels the evaluation was restricted to the proteins present in the validation dataset as they are not applicable, for proteins that don’t carry a known motif. For the motif-level enrichment, due to the possibility for partially correct hits, we used a re-implemented version of the evaluation protocol proposed in (Prytuliak et al. 2017) to compute common performance metrics (recall, precision F1, etc) both residue-wise and site-wise. Given the skewed imbalance towards higher “false positives” compared to “false negatives”, we use the

F0.5 score instead of the F1 score, which puts more weight on the precision than the recall and is calculated as follows:

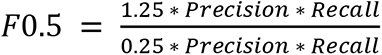

Finally, for evaluating protein-domain interactions we measured the enrichment of true-positive interactions between a given motif-carrying protein and its associated domains as reported in the validation dataset (Kumar et al. 2020). Here, true-positives represent the number of correctly associated domains for a given motif-carrying protein and then summed over all motif-carrying proteins in the predicted dataset.

### Prediction of viral SLiM binding on identified human domains

We used PepSite2 (Petsalaki et al., 2009; Trabuco et al., 2012) to predict the binding interface of each SLiM on the respective domains. Note that the PepSite2 tool tends to have a high number of false negatives but a high accuracy. PDB structures for the respective domains were selected based on the protein interaction partners for a given motif-carrying protein harboring the specified domain according to the pdb_pfam data downloaded from (http://ftp.ebi.ac.uk/pub/databases/Pfam/releases/Pfam33.1/database_files/pdb_pfamA_reg.txt.gz). The lowest resolution structure was selected if there was more than one potential structure for each domain protein. Only hits with p-value < 0.1 were considered to indicate a significant motif-domain interaction. We also applied multiple testing correction for each peptide length (3 - 10) using the Benjamini-Hochberg approach (Benjamini and Hochberg, 1995) and the same hits were retained after selecting those with adjusted p-value < 0.1. From the total of 501,109 peptide-domain pairs we identified, we were able to find an available structure for 439,192, of which 284,707 were associated with a domain protein that directly interacts with the motif-carrying protein. Those pairs were sent to PepSite2, and 76,399 peptide-domain pairs were returned as significant hits (p-value < 0.1). Since PepSite2 takes as input PDB_ID, chain and peptide sequence, some of the hits’ binding sites might not be within the specified domain region, so we filtered only those hits for a total of 46,325 peptide-domain pairs with the correct motif-domain binding interface.

### Identification of SLiM binding domains in H1 proteins

We used the iELM method (Weatheritt et al. 2012) to identify SLiM binding domains using both iELM and Pfam HMMs which are trained to recognize SLiM binding interfaces (Weatheritt et al., 2012). The HMMs were downloaded from http://elmint.embl.de/program_file/ and we used *hmmsearch* from the HMMER3 toolkit (http://hmmer.org/) to search each HMM profile (both iELM and pfam HMMs) against all H1 proteins that are reported to bind to a predicated motif instance and a viral protein. In concordance with the iELM method, we also used an E-value cut-off of 0.01 and excluded all hits with a length of <80% of the annotated SLiM-binding domain’s length. The returned domtblout output was parsed using the R package “rhmmer” (Zebulun, 2017) and ELM classes were updated to the current version of ELM database (Kumar et al., 2020) using renamed ELM classes table downloaded from ELM database (http://elm.eu.org/infos/browse_renamed.tsv).

### Clinvar mapping

Data for mapping ClinVar (Landrum et al., 2018) variants to motif residues were downloaded from (https://ftp.ncbi.nlm.nih.gov/pub/clinvar/tab_delimited/archive/submission_summary_2021-03.txt.gz) on 03/04/2021. For each motif in a given protein, we check the amino acid change of each reported variant in the corresponding protein in ClinVar and we establish a mapping if there’s at least one overlapping residue. Some of the variants in ClinVar had only genomic information with no reported amino acid changes. In that case we used the EMBL-EBI Proteins API (https://www.ebi.ac.uk/proteins/api/doc/) to retrieve the genomic coordinates of each motif that can be directly compared to those ClinVar variants with no information about amino acid changes. The mapped variants were filtered to include only single nucleotide variants (SNVs) and deletions as opposed to insertions, duplications and CNVs, since we only consider variants potentially having deleterious effects on motif function and binding affinity.

### Identification of potential Drug candidates

ChEMBL27 (Gaulton et al., 2017; Mendez et al., 2019) was downloaded from (ftp://ftp.ebi.ac.uk/pub/databases/chembl/ChEMBLdb/latest/) and data tables corresponding to compound, target and assay information were concatenated according to the database schema. The following filters were applied to retrieve high-confidence Drug-target records: 1) Only human single-protein targets were considered as opposed to protein-complex, tissue, cell_type, etc…. 2) Only compounds that are indicated to have a therapeutic application (Therapeutic_flag == 1) were considered as opposed to imaging agents, additives, etc … 3) Assays that are classified as Binding assays (Assay_type == “B”) and with high confidence score (confidence_score == 9) were considered to retrieve the assays where a precise molecular target is assigned. Using this curated dataset, we mapped the drugs that are reported to target H1 proteins (V1 → H1, where V is the viral protein). Additionally, we used the R packages: reactome.db, KEGG.db, and msigdbr to retrieve the pathways where H1 and H2 are involved, from Reactome, KEGG and MsigDB, respectively (Carlson, 2019; Dolgalev, 2020; Ligtenberg, 2019). Finally, we used the Experimental factor ontology (EFO) (Malone et al., 2010) to map the drugs and ClinVar diseases to disease ontology terms that can be systematically compared across the whole dataset. EFO disease ontologies corresponding to the identified drugs were directly retrieved from the Drug_indication table in ChEMBL database, while the ClinVar disease ontologies were cross-referenced with EFO using EMBL-EBI ontology xref service (OXO) (https://www.ebi.ac.uk/spot/oxo/index) which finds mapping between different ontology terms.

### Functional protein and domain enrichment analysis

To identify the functional pathways associated with our predicted motifs and proteins that are directly targeted by viral proteins, we performed a hypergeometric overrepresentation test on H1 proteins and PFAM (Mistry et al., 2021) domains found in both H1 proteins and those that are found to bind with our putative motif instances. For H1 proteins, we performed the enrichment against GO (biological process) (Ashburner et al., 2000; Gene Ontology Consortium, 2021) using only the domain proteins (H1) as background. We used the “enrichGO” function in the R package ClusterProfiler (Yu et al., 2012) to perform GO enrichment using default parameters except the universe parameter where we selected all 517 H1 proteins as background. We applied the enrichment on H1 proteins for all the combinations of filtered datasets and considered enriched terms with an adjusted p-value cutoff of 0.05.

Regarding domain enrichment, we used the R package “dcGOR” (Fang, 2014) to perform PFAM domain enrichment against GOBP (biological process) and using pfam2go (Mitchell et al., 2015) as background annotation which was downloaded from (http://current.geneontology.org/ontology/external2go/pfam2go) on 20.5.2021. For each set of filter combinations (Enriched Domain, PepSite, ClinVar, iELM) we used as input either the domains found in H1 proteins of the corresponding motif instances or the domains that are found in the interaction partners of the predicted motif-carrying proteins. For filter sets including “PepSite” or “Enriched Domain” we additionally required evidence of binding of the query domain to predicted motif instance or enrichment in at least 5 interacting partners of motif-carrying proteins with adjusted p-value < 0.05, respectively. Moreover, the background for each filter set was adjusted based on the initial query of the respective filter combination. For example, the background for sets including the PepSite filter would be the domains found in the original requests sent to PepSite for the respective motif instances involved. The function “dcEnrichment” was used to perform the enrichment of PFAM domains as input against their respective background. The default parameters were used except “ontology.algorithm” where we used “lea” algorithm to account for the hierarchy of the GO ontology. Specifically, once domains are already annotated to any children terms with more significance than itself, then all these domains are eliminated from the use for the recalculation of the significance at that term and the final p-values takes the maximum of the original p-value and the recalculated p-value. Finally, in order to compare the similarity of the enriched PFAM domains, we used the R package “GOSemSim” (Yu et al., 2010) to calculate the semantic similarity between the GOBP terms between each pair of PFAM domains using the “Wang” method (Wang et al., 2007). Visualization of enrichment output of “dcGOR” was implemented using the R package “ClusterProfiler” and heatmap visualization of semantic similarity was implemented using the “pheatmap” R package (Kolde, 2012).

### Enrichment of Domain-Motif interactions

Given the high false positive rate in our predicted hits and the inherent challenges in de-novo SLiM discovery, we wanted to assess whether the predicted domain-motif interactions (DMIs) are merely due to chance or can be used to infer functional SLiMs. For this purpose, we used SLiM-Enrich (Idrees et al., 2018), which calculates the enrichment of DMIs in a given PPI (protein-protein interaction) network using a permutation approach to create a background distribution of expected DMIs. We used the SLiM-Enrich Shiny webserver (http://shiny.slimsuite.unsw.edu.au/SLiMEnrich/) which takes as input PPI data in terms of domain and motif proteins. We used the ELMc-Domain strategy and either ELM as the default background for motifs or our own predicted hits from QSLiMFinder (Palopoli et al., 2015) as proxy for the motif compositions of the input motif proteins. We performed this analysis using both host-viral and human-only predicted hits in order to compare the enrichment of DMIs in both datasets.

### Context Essentiality of H1 and Motif proteins

We used CEN-tools (Sharma et al., 2020) in order to assess the essentiality of the host-proteins that are directly targeted by viral proteins and the motif-carrying proteins in our predicted motif instances. We used the classifications already provided by CEN-tools which are integrated from both Project Score (Behan et al., 2019) and DepMap (GTEx Consortium et al., 2017; Tsherniak et al., 2017) large-scale CRISPR screens in addition to a gold standard gene set from BAGEL (Hart and Moffat, 2016). We grouped both context-essential and rare-context essential so that each protein can be classified as essential, context-essential, or non-essential. Finally, we used a one-tailed Fisher’s exact test (Fisher, 1934) to perform overrepresentation of either H1 or motif-carrying proteins for each essentiality cluster in all filter combinations. Out of 517 and 2,457 H1 proteins and predicted motif proteins, 487 and 2,261 had essentiality information available, respectively and were considered for downstream analysis.

### Identification of pathway-specific domains

We used a network-based scoring approach to quantify domain-pathway associations and their specificity (Shim et al., 2019) in human domains that are found in host-viral PPI. The approach is mainly based on first constructing a domain-based network using weighted mutual information between domain profiles of human proteins, then using a Bayesian framework to assign log-likelihood scores to links of co-pathway networks. Finally, the Gini index (GI) was used to account for the distribution of each domain across pathways which eventually yields a pathway specificity (PS) score for each domain-pathway association (Shim et al., 2019). We already used the provided InterPro (Blum et al., 2021) domain profile for a total of 17,013 human proteins and 8,362 InterPro domains on both GOBP (which was already provided) and KEGG pathways (Kanehisa et al., 2021; Kanehisa and Goto, 2000) which were retrieved using the KEGGREST R package (Tenenbaum, 2020). We mapped the corresponding PFAM (Mistry et al., 2021) domains according to InterPro-PFAM cross-reference which was downloaded from (https://www.ebi.ac.uk/interpro/entry/pfam/#table), resulting in a total of 4,283 PFAM domains mapped to 4,140 out of 8,362 InterPro domains. Then, for all filter combinations, we mapped all domain-pathway associations for both KEGG and GOBP against domains in H1 proteins and domain-proteins binding to predicted motif instances. Finally, we combined all the associations and scaled their pathway-specificity scores and selected only associations with z-scores > 1 which corresponds to pathway specificity scores > 0.2.

## Notes

https://www.ebi.ac.uk/biostudies/studies/S-BSST668/)

https://gitlab.ebi.ac.uk/petsalakilab/HVSlimPred

### Competing Interest Statement

The authors have declared no competing interest.

