## Supplementary figures for "Use of viral motif mimicry improves the proteome-wide discovery of human linear motifs"

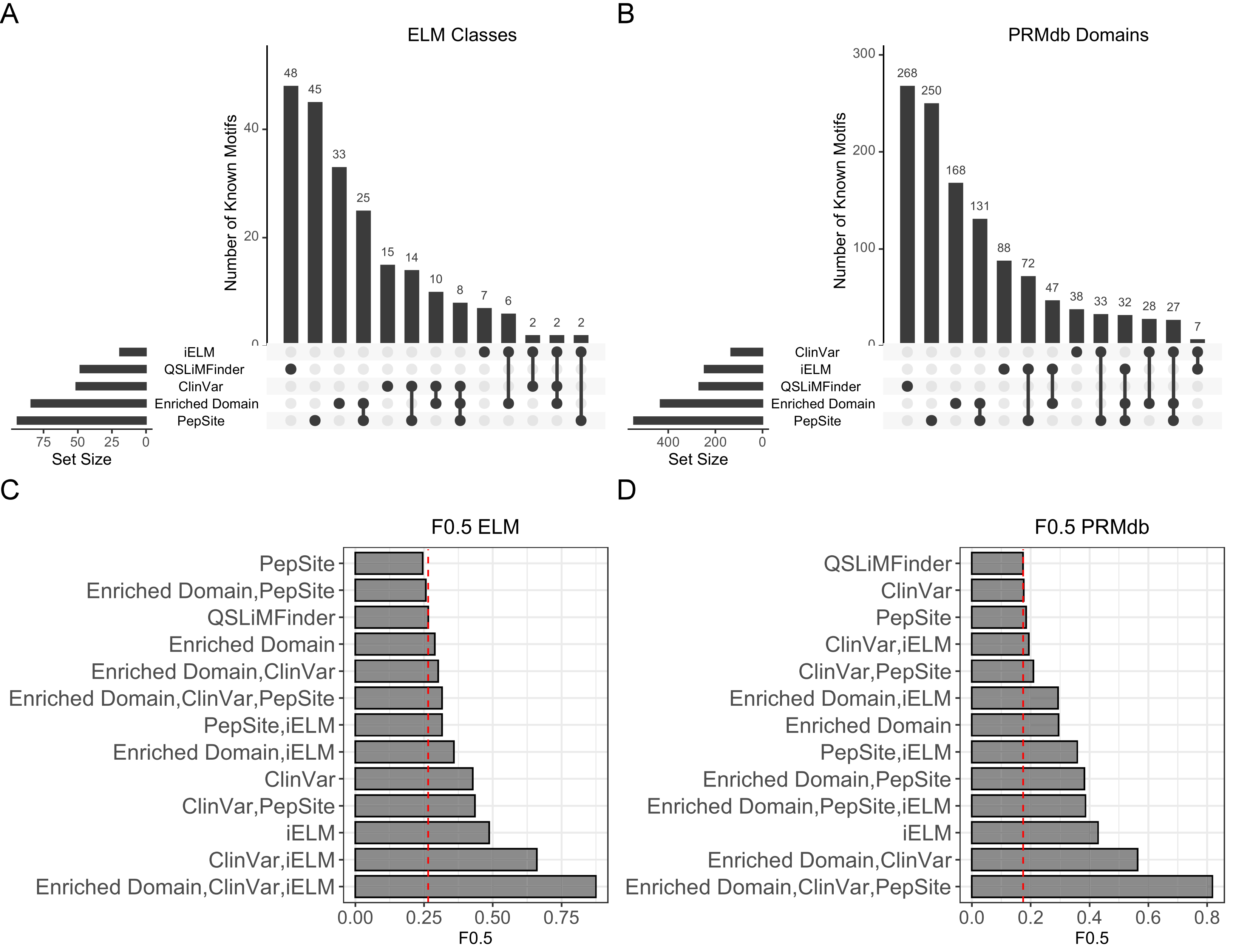
A
ELM Classes
Number of Known Motifs
48
45
33
25
15
14
10
8
7
6
2
2
2
iELM
QSLiMFinder
ClinVar
Enriched Domain
PepSite
Set Size
B
PRMdb Domains
268
250
168
131
88
72
47
38
33
32
28
27
7
C
F0.5 ELM
PepSite
Enriched Domain,PepSite
QSLiMFinder
Enriched Domain
Enriched Domain,ClinVar
Enriched Domain,ClinVar,PepSite
PepSite,iELM
Enriched Domain,iELM
ClinVar
ClinVar,PepSite
iELM
ClinVar,iELM
Enriched Domain,ClinVar,iELM
F0.5
D
F0.5 PRMdb
QSLiMFinder
ClinVar
PepSite
ClinVar,iELM
ClinVar,PepSite
Enriched Domain,iELM
Enriched Domain
PepSite,iELM
Enriched Domain,PepSite
Enriched Domain,PepSite,iELM
iELM
Enriched Domain,ClinVar
Enriched Domain,ClinVar,PepSite

**Figure S1: Evaluation of known motif instances.** (A & B) UpSet plot showing the overlapping number of known ELM classes (A) or PRMdb domains (B) and predicted datasets defined by a single or multiple filters. (C & D) Bar graph plotting the enrichment of known ELM classes (C) or PRMdb domains (D) represented by F0.5 score for each set of applied filters. (See also **Figure 2**)

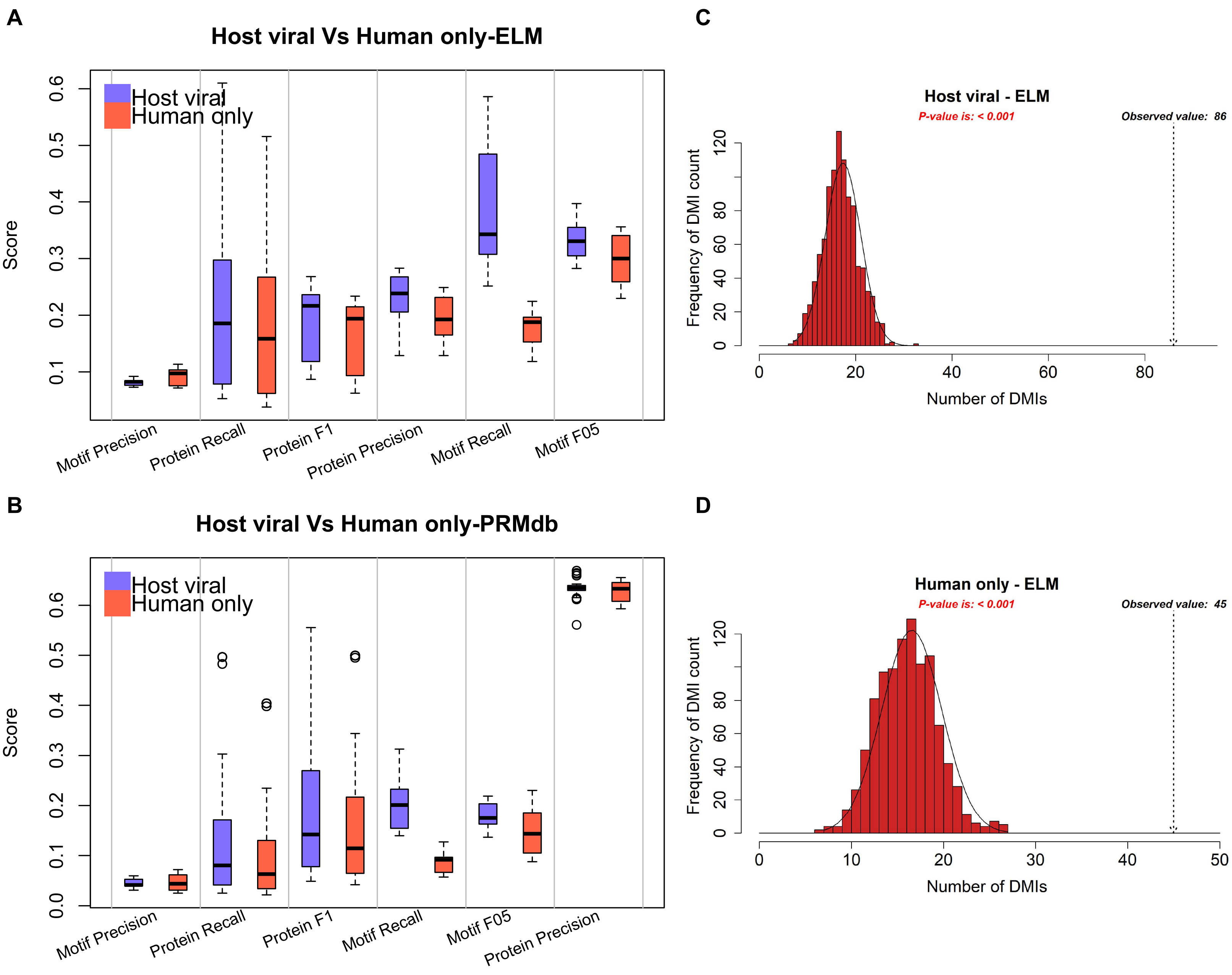
A
Host viral Vs Human only-ELM
Host viral
Human only
Score
Motif Precision
Protein Recall
Protein F1
Protein Precision
Motif Recall
Motif F05
B
Host viral Vs Human only-PRMdb
Host viral
Human only
Score
Motif Precision
Protein Recall
Protein F1
Motif Recall
Motif F05
Protein Precision
C
Host viral - ELM
P-value is: < 0.001
Observed value: 86
Frequency of DMI count
Number of DMIs
D
Human only - ELM
P-value is: < 0.001
Observed value: 45
Frequency of DMI count
Number of DMIs

**Figure S2: Comparison of known motifs and domain-motif interactions between Host-viral and Human-only predictions.** (A & B) Box plot showing the distribution of performance metrics for all 60 datasets created by different combinations of parameters defined by enriched domains in the interacting partners of motif-carrying proteins (**Methods**), in addition to the raw predicted output from QSLiMFinder. The metrics were calculated using known ELM classes (A) and PRMdb domains (B) as background, respectively. Precision, recall and F score were calculated based on evaluation on the motif-level and protein level (**Methods**)., F05, Fbeta measure (beta = 0.5); Motif, Motif-level evaluation; Protein, Protein-level evaluation. (C & D) Histogram showing the expected distribution of predicted domain-motif interactions (DMIs) from 1000 randomized PPI data from host-viral (C) and human only (D) predicted instances. The observed number of non-redundant predicted DMIs in the real data is also marked.

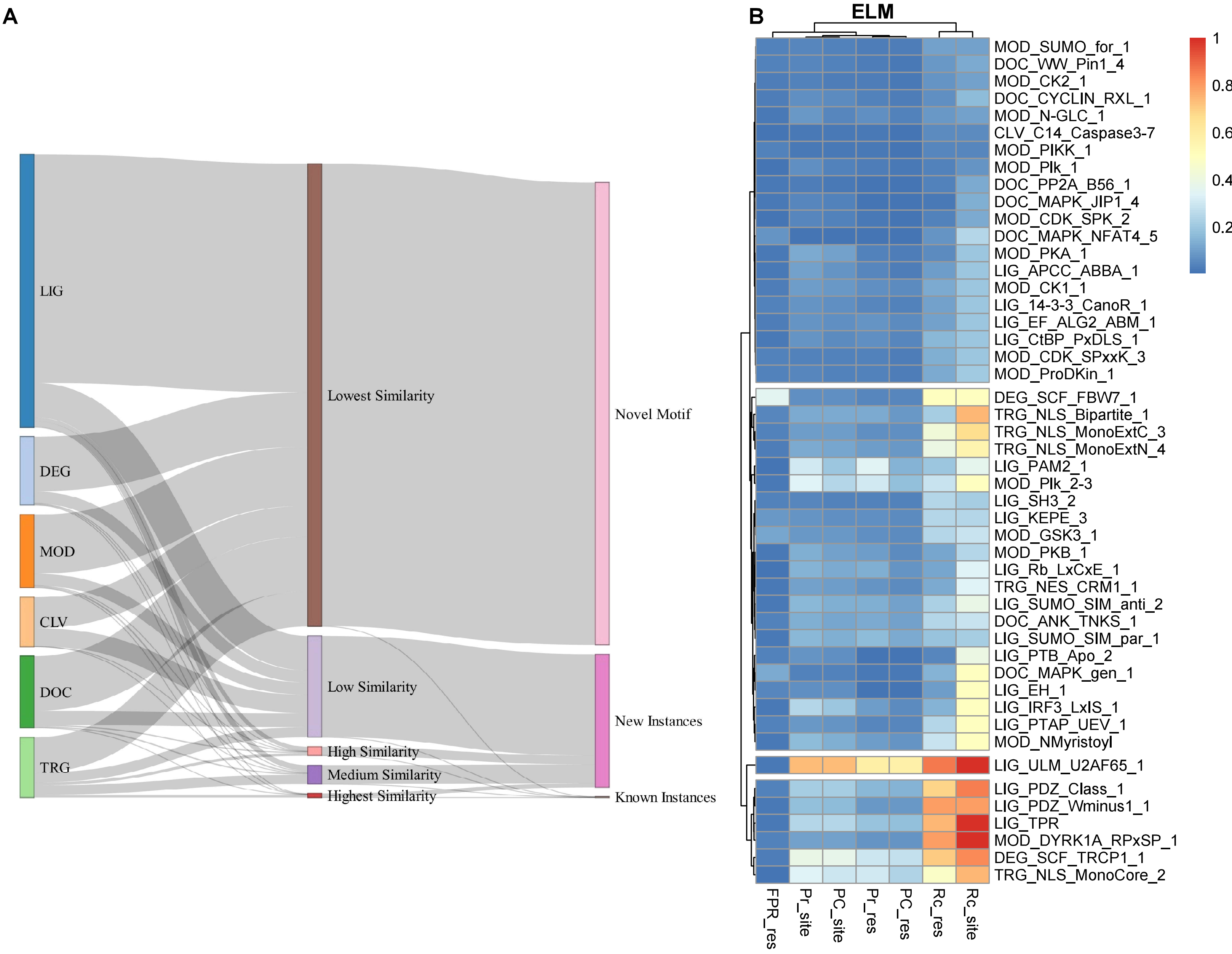
A
LIG
DEG
MOD
CLV
DOC
TRG
Lowest Similarity
Low Similarity
High Similarity
Medium Similarity
Highest Similarity
Novel Motif
New Instances
Known Instances
B
ELM
MOD_SUMO_for_1
DOC_WW_Pin1_4
MOD_CK2_1
DOC_CYCLIN_RXL_1
MOD_N-GLC_1
CLV_C14_Caspase3-7
MOD_PIKK_1
MOD_Plk_1
DOC_PP2A_B56_1
DOC_MAPK_JIP1_4
MOD_CDK_SPK_2
DOC_MAPK_NFAT4_5
MOD_PKA_1
LIG_APCC_ABBA_1
MOD_CK1_1
LIG_14-3-3_CanoR_1
LIG_EF_ALG2_ABM_1
LIG_CtBP_PxDLS_1
MOD_CDK_SPxxK_3
MOD_ProDKin_1
DEG_SCF_FBW7_1
TRG_NLS_Bipartite_1
TRG_NLS_MonoExtC_3
TRG_NLS_MonoExtN_4
LIG_PAM2_1
MOD_Plk_2-3
LIG_SH3_2
LIG_KEPE_3
MOD_GSK3_1
MOD_PKB_1
LIG_Rb_LxCxE_1
TRG_NES_CRM1_1
LIG_SUMO_SIM_anti_2
DOC_ANK_TNKS_1
LIG_SUMO_SIM_par_1
LIG_PTB_Apo_2
DOC_MAPK_gen_1
LIG_EH_1
LIG_IRF3_LxIS_1
LIG_PTAP_UEV_1
MOD_NMyristoyl
LIG_ULM_U2AF65_1
LIG_PDZ_Class_1
LIG_PDZ_Wminus1_1
LIG_TPR
MOD_DYRK1A_RPxSP_1
DEG_SCF_TRCP1_1
TRG_NLS_MonoCore_2
FPR_res
Pr_site
PC_site
Pr_res
PC_res
Rc_res
Rc_site
1
0.8
0.6
0.4
0.2

**Figure S3: Classification and comparison of predicted instances to known ELM classes** (A) Sankey network for drawing the similarity between the predicted instances and their corresponding ELM classes based on their CompairMotif similarity score. The similarity scores were divided into 5 levels ranked from Highest to Lowest according to the following thresholds: Highest, >= 70%; High, >= 60%; Medium, >= 50%; Low, >= 40%; Lowest, < 40%. Predicted instances were classified as known if the predicted motif was found in the right protein and the right location compared to known ELM instances, while they were classified as new or novel instances if the predicted motif pattern was higher or lower than 40% similarity compared to a given ELM class, respectively. LIG,ligand; CLV,cleavage; DOC,docking; DEG,degradation; MOD,post-translational modification; TRG,targeting. (B) Heatmap showing performance metrics for known ELM classes that are recovered by at least one predicted instance (i.e sharing common motif residues). The metrics were calculated at the residue-level, comparing predicted and known motif residues and at the site-level where a site is defined by at least one common residue between the predicted and ELM motif in each instance. Pr, precision; Rc, Recall; FPR, False positive rate.

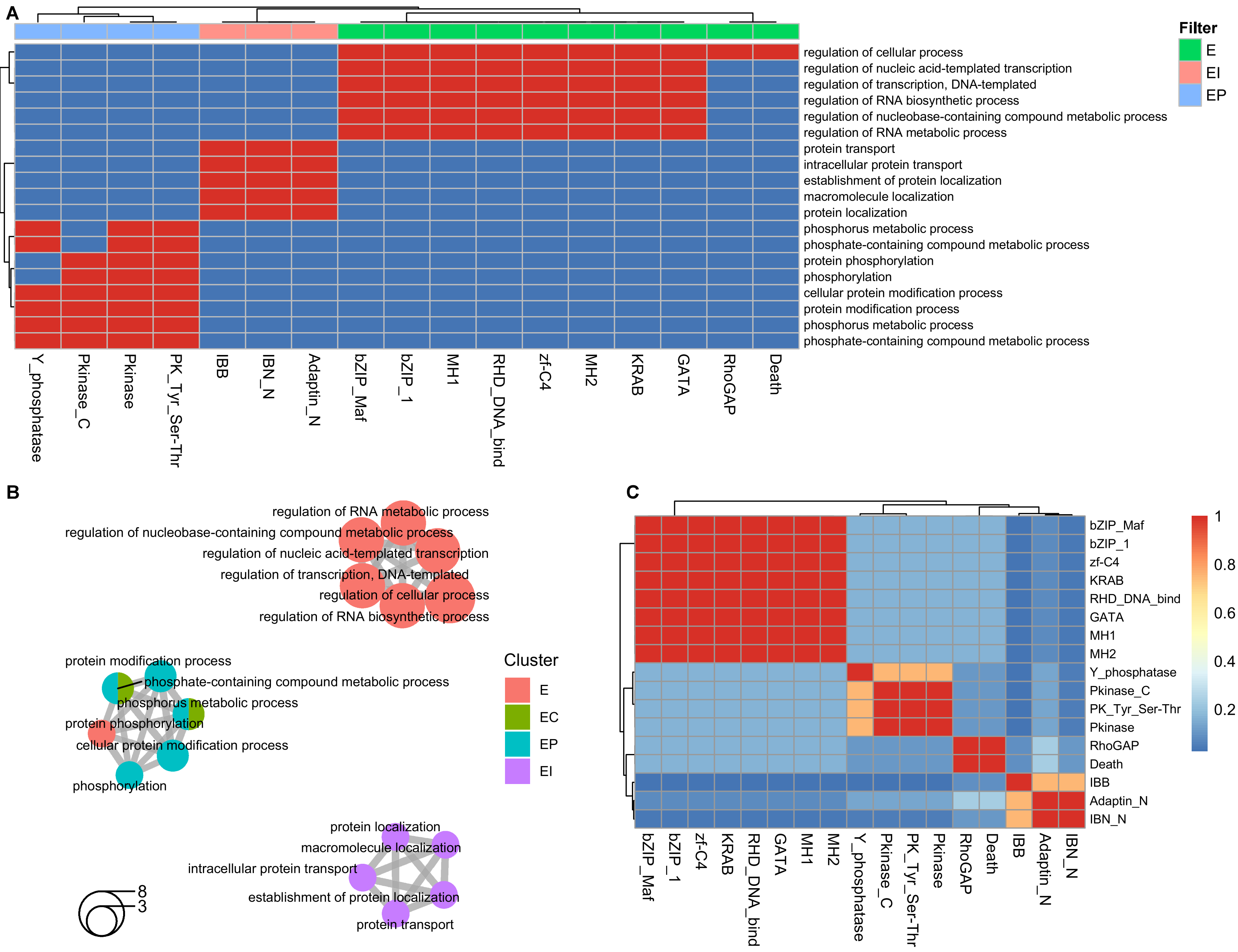
A
Filter
E
EI
EP
regulation of cellular process
regulation of nucleic acid-templated transcription
regulation of transcription, DNA-templated
regulation of RNA biosynthetic process
regulation of nucleobase-containing compound metabolic process
regulation of RNA metabolic process
protein transport
intracellular protein transport
establishment of protein localization
macromolecule localization
protein localization
phosphorus metabolic process
phosphate-containing compound metabolic process
protein phosphorylation
phosphorylation
cellular protein modification process
protein modification process
phosphorus metabolic process
phosphate-containing compound metabolic process
Y_phosphatase
Pkinase_C
Pkinase
PK_Tyr_Ser-Thr
IBB
IBN_N
Adaptin_N
bZIP_Maf
bZIP_1
MH1
RHD_DNA_bind
zf-C4
MH2
KRAB
GATA
RhoGAP
Death
B
regulation of RNA metabolic process
regulation of nucleobase-containing compound metabolic process
regulation of nucleic acid-templated transcription
regulation of transcription, DNA-templated
regulation of cellular process
regulation of RNA biosynthetic process
protein modification process
phosphate-containing compound metabolic process
phosphorus metabolic process
protein phosphorylation
cellular protein modification process
phosphorylation
Cluster
E
EC
EP
EI
protein localization
macromolecule localization
intracellular protein transport
establishment of protein localization
protein transport
8
3
C
bZIP_Maf
bZIP_1
zf-C4
KRAB
RHD_DNA_bind
GATA
MH1
MH2
Y_phosphatase
Pkinase_C
PK_Tyr_Ser-Thr
Pkinase
RhoGAP
Death
IBB
Adaptin_N
IBN_N
1
0.8
0.6
0.4
0.2

**Figure S4: GO pathway enrichment of PFAM domains in interacting partners of motif-carrying proteins** (A) Binary heatmap describing association between enriched GO terms (biological process) and the corresponding overlapping domains between input dataset and PFAM2GOBP annotations for a given term, where red and blue boxes represent presence or absence of enriched terms for a given domain, respectively. Only Domain-pathway pairs with adjusted p-value < 0.05 are displayed. Domains are clustered by filter combinations shown as colored annotation bars above columns.E, Enriched Domains; P, PepSite; C,ClinVar; I, iELM. (B) Network showing similarity between enriched GO terms which are calculated according to Jaccard similarity coefficient. Each node is a pie chart colored by filter combinations and the size of the nodes correspond to the number of domains. E, Enriched Domains; P, PepSite; C,ClinVar; I, iELM. (C) Heatmap showing GO semantic similarity between enriched domains calculated using the “Wang" method (Wang et al., 2007) between GO terms for each domain-pair. The color scale represents the similarity score where 1 denotes complete overlap, and 0 denotes no similarity.

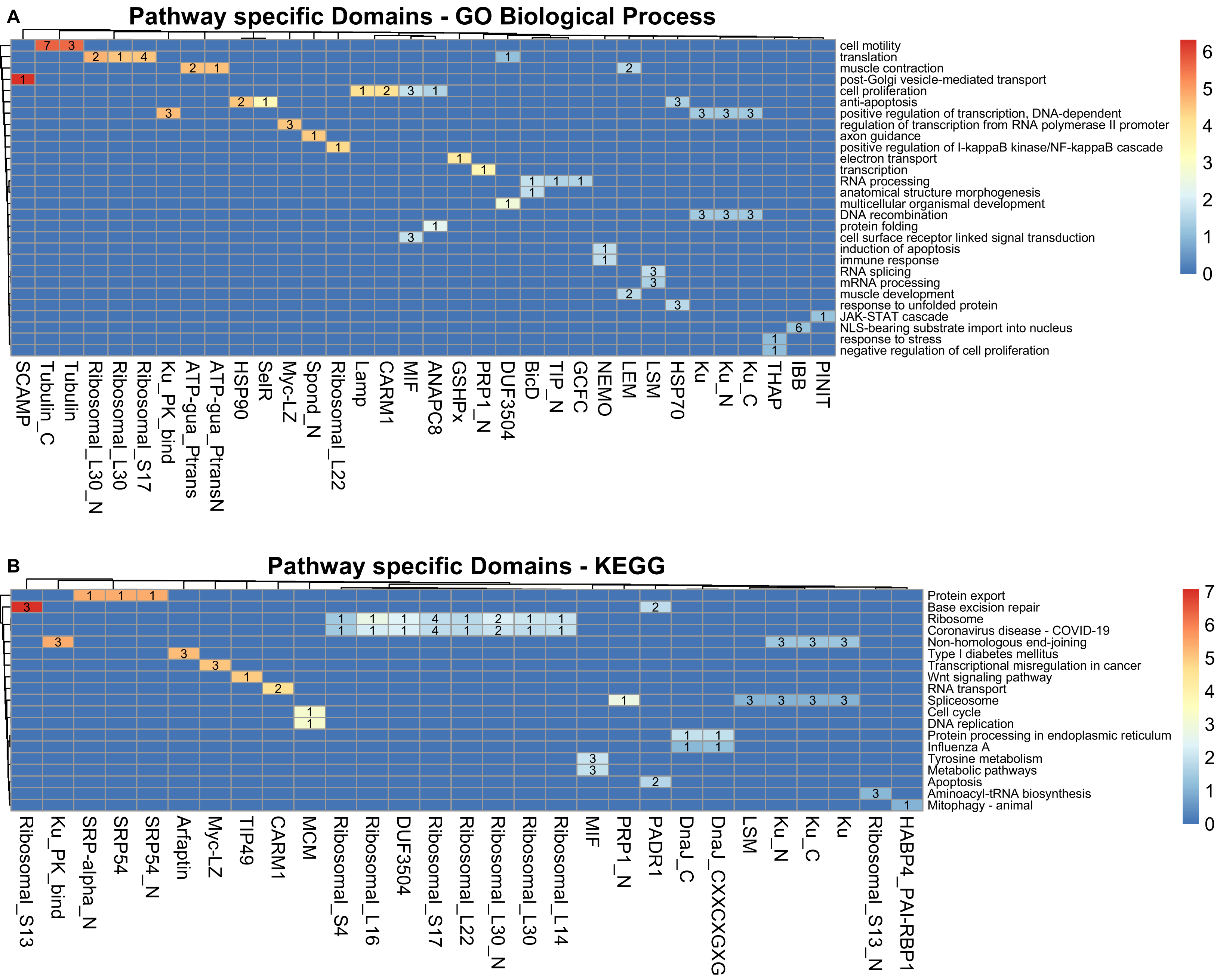
A
Pathway specific Domains - GO Biological Process
cell motility
translation
muscle contraction
post-Golgi vesicle-mediated transport
cell proliferation
anti-apoptosis
positive regulation of transcription, DNA-dependent
regulation of transcription from RNA polymerase II promoter
axon guidance
positive regulation of I-kappaB kinase/NF-kappaB cascade
electron transport
transcription
RNA processing
anatomical structure morphogenesis
multicellular organismal development
DNA recombination
protein folding
cell surface receptor linked signal transduction
induction of apoptosis
immune response
RNA splicing
mRNA processing
muscle development
response to unfolded protein
JAK-STAT cascade
NLS-bearing substrate import into nucleus
response to stress
negative regulation of cell proliferation
SCAMP
Tubulin_C
Tubulin
Ribosomal_L30_N
Ribosomal_L30
Ribosomal_S17
Ku_PK_bind
ATP-gua_Ptrans
ATP-gua_PtransN
HSP90
SelR
Myc-LZ
Spond_N
Ribosomal_L22
Lamp
CARM1
MIF
ANAPC8
GSHPx
PRP1_N
DUF3504
BicD
TIP_N
GCFC
NEMO
LEM
LSM
HSP70
Ku
Ku_N
Ku_C
THAP
IBB
PINIT
B
Pathway specific Domains - KEGG
Protein export
Base excision repair
Ribosome
Coronavirus disease - COVID-19
Non-homologous end-joining
Type I diabetes mellitus
Transcriptional misregulation in cancer
Wnt signaling pathway
RNA transport
Spliceosome
Cell cycle
DNA replication
Protein processing in endoplasmic reticulum
Influenza A
Tyrosine metabolism
Metabolic pathways
Apoptosis
Aminoacyl-tRNA biosynthesis
Mitophagy - animal
Ribosomal_S13
Ku_PK_bind
SRP-alpha_N
SRP54
SRP54_N
Arfaptin
Myc-LZ
TIP49
CARM1
MCM
Ribosomal_S4
Ribosomal_L16
DUF3504
Ribosomal_S17
Ribosomal_L22
Ribosomal_L30_N
Ribosomal_L30
Ribosomal_L14
MIF
PRP1_N
PADR1
DnaJ_C
DnaJ_CXXCXGXG
LSM
Ku_N
Ku_C
Ku
Ribosomal_S13_N
HABP4_PAI-RBP1

**Figure S5: Pathway-specific domains in H1 proteins** (A & B) Heatmap showing the relationship between pathway-specific domains and their associated pathways using either GO (A) or KEGG (B) as background. The color scale represents z-score standardization of pathway-specificity scores (i.e confidence score) for each domain-pathway association (DPA). Only DPAs with z-score > 1 are shown and labelled according to the number of filter combinations having the respective domain in at least one of the H1 proteins involved.

**A**

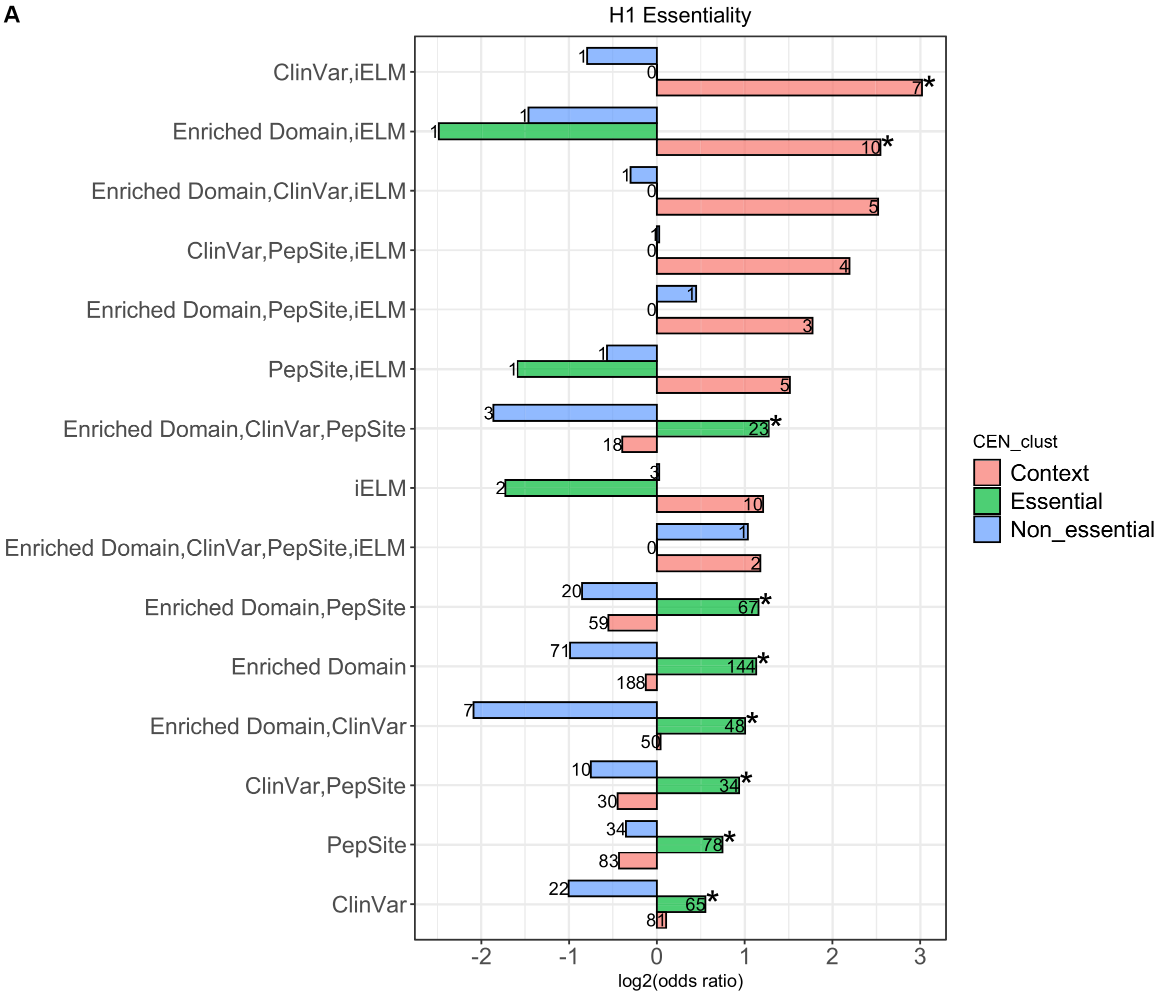
H1 Essentiality
ClinVar,iELM
Enriched Domain,iELM
Enriched Domain,ClinVar,iELM
ClinVar,PepSite,iELM
Enriched Domain,PepSite,iELM
PepSite,iELM
Enriched Domain,ClinVar,PepSite
iELM
Enriched Domain,ClinVar,PepSite,iELM
Enriched Domain,PepSite
Enriched Domain
Enriched Domain,ClinVar
ClinVar,PepSite
PepSite
ClinVar
log2(odds ratio)
CEN_clust
Context
Essential
Non_essential

**B**

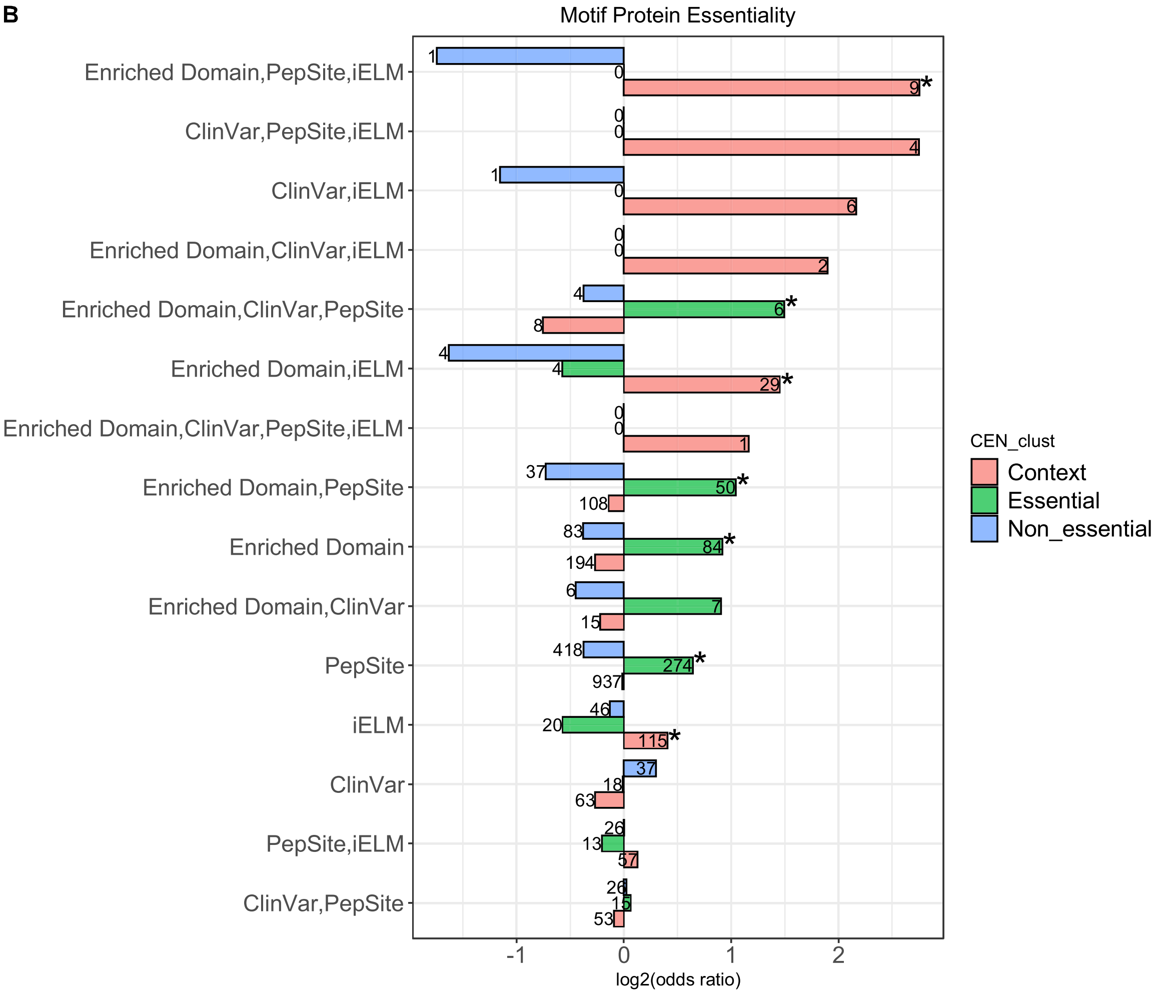
Motif Protein Essentiality
Enriched Domain,PepSite,iELM
ClinVar,PepSite,iELM
ClinVar,iELM
Enriched Domain,ClinVar,iELM
Enriched Domain,ClinVar,PepSite
Enriched Domain,iELM
Enriched Domain,ClinVar,PepSite,iELM
Enriched Domain,PepSite
Enriched Domain
Enriched Domain,ClinVar
PepSite
iELM
ClinVar
PepSite,iELM
ClinVar,PepSite
log2(odds ratio)
CEN_clust
Context
Essential
Non_essential

**Figure S6: Context essentiality of H1 and motif proteins** (A & B) Bar graph plotting the enrichment of H1 (A) and motif (B) proteins for each of the essentiality classification as determined by CEN-tools (Methods) in each set of applied filters. Odds ratio was calculated based on a one-tailed fisher-exact test using H1 proteins (A) or motif protein predicted by QSLiMFinder (B) as background. The bars are colored by the essentiality classification (context, essential and non-essential) and labelled by the number of query proteins in each dataset. Asterisks are only added to denote significant enrichment according to the fisher-exact test ($p$-value $< 0.05$).

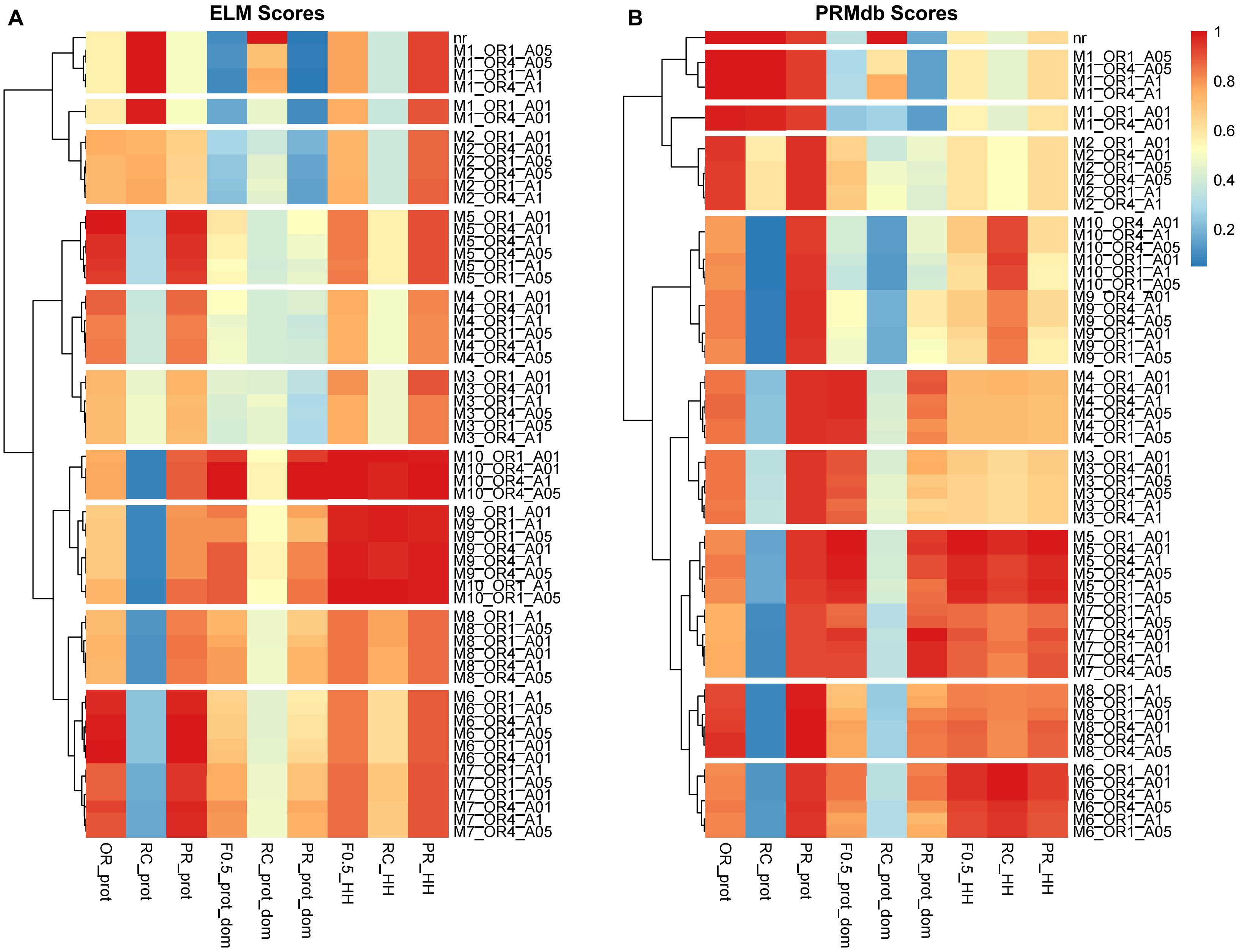
A
ELM Scores
nr
M1_OR1_A05
M1_OR4_A05
M1_OR1_A1
M1_OR4_A1
M1_OR1_A01
M1_OR4_A01
M2_OR1_A01
M2_OR4_A01
M2_OR1_A05
M2_OR4_A05
M2_OR1_A1
M2_OR4_A1
M5_OR1_A01
M5_OR4_A01
M5_OR4_A1
M5_OR4_A05
M5_OR1_A1
M5_OR1_A05
M4_OR1_A01
M4_OR4_A01
M4_OR1_A1
M4_OR1_A05
M4_OR4_A1
M4_OR4_A05
M3_OR1_A01
M3_OR4_A01
M3_OR1_A1
M3_OR4_A05
M3_OR1_A05
M3_OR4_A1
M10_OR1_A01
M10_OR4_A01
M10_OR4_A1
M10_OR4_A05
M9_OR1_A01
M9_OR1_A1
M9_OR1_A05
M9_OR4_A01
M9_OR4_A1
M9_OR4_A05
M10_OR1_A1
M10_OR1_A05
M8_OR1_A1
M8_OR1_A05
M8_OR1_A01
M8_OR4_A01
M8_OR4_A1
M8_OR4_A05
M6_OR1_A1
M6_OR1_A05
M6_OR4_A1
M6_OR4_A05
M6_OR1_A01
M6_OR4_A01
M7_OR1_A1
M7_OR1_A05
M7_OR1_A01
M7_OR4_A01
M7_OR4_A1
M7_OR4_A05
OR_prot
RC_prot
PR_prot
F0.5_prot_dom
RC_prot_dom
PR_prot_dom
F0.5_HH
RC_HH
PR_HH
B
PRMdb Scores
nr
M1_OR1_A05
M1_OR4_A05
M1_OR1_A1
M1_OR4_A1
M1_OR1_A01
M1_OR4_A01
M2_OR1_A01
M2_OR4_A01
M2_OR1_A05
M2_OR4_A05
M2_OR1_A1
M2_OR4_A1
M10_OR4_A01
M10_OR4_A1
M10_OR4_A05
M10_OR1_A01
M10_OR1_A1
M10_OR1_A05
M9_OR4_A01
M9_OR4_A1
M9_OR4_A05
M9_OR1_A01
M9_OR1_A1
M9_OR1_A05
M4_OR1_A01
M4_OR4_A01
M4_OR4_A1
M4_OR4_A05
M4_OR1_A1
M4_OR1_A05
M3_OR1_A01
M3_OR4_A01
M3_OR1_A05
M3_OR4_A05
M3_OR1_A1
M3_OR4_A1
M5_OR1_A01
M5_OR4_A01
M5_OR4_A1
M5_OR4_A05
M5_OR1_A1
M5_OR1_A05
M7_OR1_A1
M7_OR1_A05
M7_OR4_A01
M7_OR1_A01
M7_OR4_A1
M7_OR4_A05
M8_OR1_A1
M8_OR1_A05
M8_OR1_A01
M8_OR4_A01
M8_OR4_A1
M8_OR4_A05
M6_OR1_A01
M6_OR4_A01
M6_OR4_A1
M6_OR4_A05
M6_OR1_A1
M6_OR1_A05
OR_prot
RC_prot
PR_prot
F0.5_prot_dom
RC_prot_dom
PR_prot_dom
F0.5_HH
RC_HH
PR_HH
1
0.8
0.6
0.4
0.2

**Figure S7: Evaluation of known ELM instances in different combinations of enriched domains' datasets** (A & B) Heatmap showing normalized performance metrics for known ELM instances in different datasets built from a combination of 3 parameters to define the enriched domain filter: Minimum number of interacting partners bearing the enriched domain (denoted by M), minimum odds ratio as determined by a one-tailed fisher-exact test (denoted by OR), and adjusted p-value cutoff (denoted by A). Each metric (column) is divided by their maximum value to obtain 0-1 normalized values for each dataset. Precision, recall and F score were calculated based on evaluation at the motif-level, protein level, and protein-domain level, while odds ratio was calculated only at the protein-level (Methods). PR, precision; RC, Recall; F0.5, Fbeta measure (beta = 0.5); HH, Motif-level evaluation; prot, Protein-level evaluation; prot_dom, Protein-domain level evaluation; OR, odds-ratio.
